# Long-range genomic loci stochastically assemble into hierarchical chromosome structures

**DOI:** 10.1101/2025.02.10.637328

**Authors:** Jingyu Zhang, Siyuan Wang, Simon C Watkins, Jianhua Xing

**Affiliations:** Department of Computational and Systems Biology, University of Pittsburgh; Pittsburgh, PA 15232, USA; Department of Cell Biology, Yale School of Medicine; New Haven, CT 06510, USA; Department of Cell Biology, University of Pittsburgh; Pittsburgh, PA 15232, USA; UPMC-Hillman Cancer Center, University of Pittsburgh; Pittsburgh, PA, USA; Department of Physics and Astronomy, University of Pittsburgh; Pittsburgh, PA 15232, USA

**Keywords:** genome folding, chromatin loop organization, long-range colocalization, chromosome territories, multivalent binding

## Abstract

**Background:** One fundamental yet open question is how eukaryotic chromosomes fold into segregated territories, a process essential for gene transcription and cell fate. While several models describe how chromosomal loops form at shorter scales, the mechanisms governing large-scale (> 100 Mb) chromosome organization remain poorly understood.

**Results:** Through analyzing multiple sequencing- and imaging-based datasets, we identify long-range chromosomal backbone loop structures that span over 100 Mb, extending beyond the reach of several existing DNA loop models. Some long-range loops are stable for at least 10 minutes, as shown by live-cell imaging with sequence-specific CRISPR-dCas9 fluorescent labeling. Biophysical modeling further demonstrates that their formation is driven by a multivalent binding mechanism. Further epigenetic profiling and spatial density analyses indicate that many of the assembly formations are independent of known large-scale nuclear structures.

**Conclusions:** Our findings suggest a redundant, distributed cluster mechanism that ensures chromosomal organization robustness across cell types and against mutations. This mechanism coordinates large-scale chromosome compaction while simultaneously guiding the formation of smaller-scale chromosomal structures, offering a new framework for understanding genome folding and its role in cell identity.

## Background

A central problem in structural biology and biological physics is how macromolecules fold into three-dimensional structures. An emerging “local-to-global” mechanism highlights the role of local cooperativity among peptide monomers in guiding the formation of a native protein structure from an expansive conformational space [1]. In contrast to typical polypeptides with 300 – 400 amino acids, the normal human genome is composed of 46 long-chain polymers, ranging from ∼ 40 to 250 Mb, with a total linear length extending up to two meters. Given that 3D structures play a central role in regulating cell identity and gene expression [2], an intriguing question is how these linear DNA molecules compact into segregated chromatin territories within a cell nucleus, typically only about 10 μm in diameter.

Chromatin, unlike a polypeptide, is often considered to lack a stable, ordered three-dimensional structure due to its significantly greater length. When modeled as a linear polymer chain without structural supports such as protein bridges or long non-coding RNA networks, theoretical analyses predict that the average contact frequency (*f*) between two chromosomal fragments depends on their genomic separation (*l*) and follows a power-law as *f* ∝ *l*^α^. Previous Hi-C measurements have confirmed this relationship, with an exponent α ∼ –1 within the 1 – 5 Mb range (Fig. 1a, Supplementary Fig. S1a-c). This result aligns with polymer models of chromatin as a crumpled polymer [3–5] or that emphasize the excluded-volume effect [6].

**Fig. 1.**
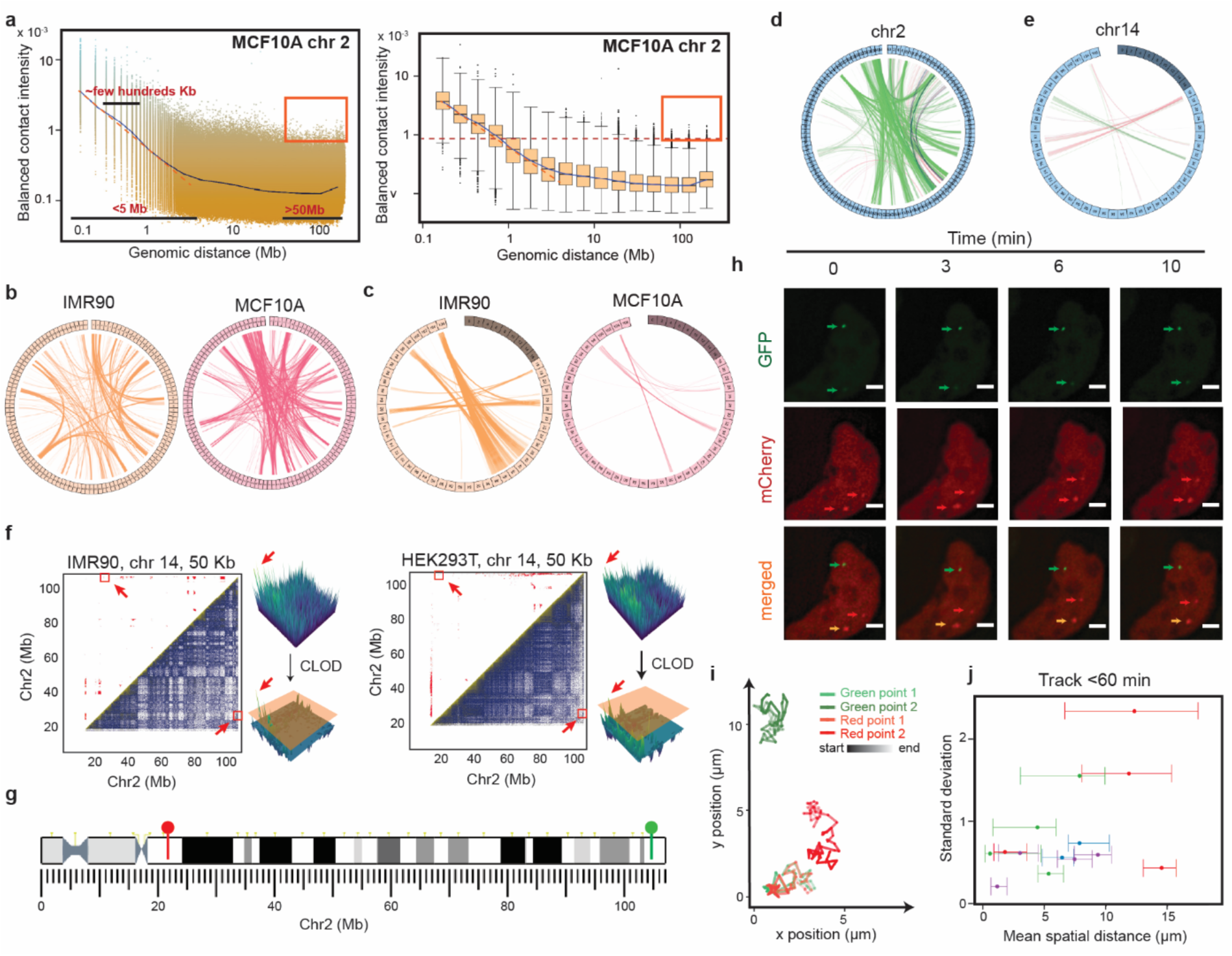
Hi-C data reveal the prevalent existence of colocalized pairs of long-range genomic loci. (**a**) Scatter plot (left) and boxplot (right) showing the decay trend (blue line) of balanced contact intensity with increasing genomic distance. Red boxed points represent locus pairs > 50 Mb apart with contact intensity exceeding the third quartile of those observed for ∼1 Mb pairs. (**b, c**) Representative linkage maps of colocalized locus pairs > 50 Mb on chr2 and chr14 in two different cell lines. (**d, e**) Linkage maps of colocalized pairs > 50 Mb apart, shared across at least two of the following cell types: four embryonic cell lines (green), four normal differentiated cell lines (pink), and four cancer cell lines (blue). (**f**) CLOD-based identification of colocalized pairs on chr14 from Hi-C data of IMR90 and HEK293T cells at 50 kb resolution. (**g**) CRISPR-dCas9 two-color labeling sites on chr14. (**h**) Live-cell imaging snapshots of a HEK293T cell labeled with RFP and GFP. Scale bar: 2 *μm*. (**i**) 10-min time-resolved trajectories of green and red puncta in the labeled cell from panel h. (**j**) Distance fluctuations between green and red puncta pairs in four cells that were tracked for 60 min. The left and right bars indicate the maximum and minimum distances between a pair of puncta.

However, deviations from the power-law behavior have been observed at both short and long genomic distances. At genomic distances ∼100 – 200 kb, the deviations (Fig. 1a, Supplementary Fig. S1a-c) suggest the presence of stable chromatin loops, potentially formed through a loop-extrusion mechanism [7]. Beyond 5 Mb, Hi-C data reveal further deviations, with the *f*-*l* curves leveling off (Fig. 1a, blue line; Supplementary Fig. S1a&b). This phenomenon has been attributed to polymer models incorporating random-sized loop formation between chromatin segments [8]. Additional factors, such as loose/compact compartment segregation and nuclear structure tethering, also contribute to deviations from the crumpled polymer model at large genomic distances [4, 9].

In sum, previous studies suggest that chromosomes form functionally important local structures such as topologically associated domains (TADs) and enhancer-promoter interactions, typically spanning within ∼ 1 Mb. Reports also exist of stable, long-range genomic interactions (>10 Mb) in large genomes of species including human [4, 9–11], and mechanisms such as liquid-liquid phase separation mediated by Polycomb and HP1 are known to facilitate some such long-range contacts [12, 13]. However, there is a lack of systematic investigations on the existence of sequence-specific, conserved, and stable locus-locus interactions >50 Mb and the physical principles governing them. Here, we address this gap by characterizing spatial colocalization of pairs of distant loci on chromosomes of several mammalian and non-mammalian species. We demonstrate the conservation of their spatial proximity across diverse cell lines and elucidate the physical mechanisms that likely enforce this stable association.

## Results

### Hi-C data reveals the prevalence of long-range genomic locus colocalization beyond 50 Mb

We first analyzed Hi-C data for chromosome 2 (chr2) and chromosome 14 (chr14) in human MCF10A cells [14]. While most genomic pairs followed the expected *f*-*l* curve, certain pairs (e.g., red box in Fig. 1a & Supplementary Fig. S1a) showed significant deviations, with contact frequencies comparable to the median value of 1-Mb pairs despite their much larger genomic separation. Similar distal locus pairs with high contact frequencies were also observed in chr2 and chr14 of human IMR90 cells [15] (Supplementary Fig. S1b). Extending this analysis to 11 additional human cell lines revealed that such deviations are prevalent among genomic locus pairs separated by even more than 50 Mb (Supplementary Fig. S2), suggesting their stable three-dimensional proximity in at least one subpopulation of chromatin. Notably, these long-range interactions extend beyond typical intrachromosomal contacts such as enhancer-promoter interactions, super-enhancers, and TADs [11, 13]. Given the entropic cost of maintaining substantial colocalization probability over such long genomic distances, we sought to explore their underlying mechanisms and potential functional significance.

While the above statistical analyses identified long-range locus pairs with spatial proximity, detecting them quantitatively from noisy Hi-C data remains challenging, as strong signals from short-range locus pairs often overshadow weaker signals from long-range ones. To address this challenge, we developed a computational pipeline, <u>C</u>hromosome <u>L</u>ong-range colocalization identifier through <u>O</u>utlier <u>D</u>etection (CLOD) (Supplementary Fig. S3 and Methods). When we performed CLOD on actual Hi-C data, it successfully identified colocalized bins that have been proven via other experiments. For example, CLOD identified a 150 kb loop (Supplementary Fig. S1d), which was previously validated as a promoter-enhancer interaction of PRIP1 using Hi-C [16], from the MCF7 Hi-C dataset [14]. We then applied CLOD to a Hi-C dataset from the HCT116 cell line [17] and detected a colocalized genomic pair separated by 27 Mb (Supplementary Fig. S1e). This locus pair was independently confirmed by fluorescence in situ hybridization (FISH) [10]. Collectively, these results demonstrate that the CLOD pipeline can effectively detect colocalized locus pairs across both short and long genomic distances.

We extended our CLOD analysis to other chromosomes longer than 100 Mb in the IMR90 cell line and found that long-range contacting pairs are prevalent in all long chromosomes (Supplementary Fig. S4a). We further applied CLOD to Hi-C datasets from a panel of four normal differentiated human cell lines, five cancer cell lines, and four embryonic cell lines [18–20] (Supplementary Table S1). Across these datasets, we identified multiple colocalized locus pairs separated by 50 Mb or more on chr2 and chr14 (Fig. 1b&c and Supplementary Fig. Sb). Notably, some of these long-range pairs were consistently detected across multiple cell lines of different types (Fig. 1d&e). However, about half (43% on chr2 and 52% on chr14) of the identified long-range pairs are cell-type-specific, and fewer than 0.1% are detected in at least half of the tested cell lines. To assess whether the colocalization is restricted to human chromosomes, we examined colocalizations in long (>150 Mb) chromosomes of other species, including six mammals (*Mus musculus* (mouse) [21, 22], *Rattus norvegicus* (rat), *Macaca mulatta* (rhesus monkey), *Bos taurus* (cattle), *Ovis aries* (sheep), *Oryctolagus cuniculus* (rabbit) and *Gallus gallus* (chicken) [22]. Consistently, we identified colocalization of a substantial number of genomic locus pairs spanning genomic distances exceeding 50 Mb (Supplementary Fig. S5, Supplementary Table S1), suggesting this spatial organization is broadly conserved.

To determine whether the CLOD-identified long-range colocalization was transient or persistent, we focused on a colocalized genomic pair on human chr14, separated by approximately 80 Mb, that had been detected in multiple cell lines, including IMR90 and HEK293T (Fig. 1f) [23]. We labeled the loci with two-color CRISPR-dCas9-guided fluorescent proteins in HEK293T cells (Fig. 1g, Methods, Supplementary Table S2). Live-cell imaging confirmed the sustained proximity and coordinated movement of green and red puncta, with their co-fluctuations persisting throughout the observed 10 min in some cases (Fig. 1h&i, and Supplementary Movie S1). Extended imaging further showed that the neighboring puncta remained in proximity in the cells examined during an imaging period of about 1 hour (Fig. 1j and Supplementary Fig. S1f).

Therefore, from Hi-C data via CLOD, we identified the prevalence of long-range colocalizations (> 50 Mb) across multiple human cell lines, including normal, cancer, and embryonic cells, as well as in diverse other species from mammals to chicken. Such distal genomic colocalizations may be temporally persistent, as confirmed by live-cell imaging.

### MERFISH chromatin tracing data reveal prevalence and conservation of colocalized loci > 100 Mb in human chr2

While bulk Hi-C provides an ensemble-averaged view of chromosome conformations, it lacks the resolution to capture individual chromosome structures. It can neither distinguish whether colocalized pairs originate from the same or different homologous chromosomes, nor determine whether the observed contact frequencies reflect widespread interactions among cells or rare subpopulations. To cross-validate the prevalence of long-range pair colocalization and assess their distribution at the single-chromatin level, we analyzed previously published IMR90 chromatin tracing data obtained using DNA MERFISH [24].

As a technical control, we first re-analyzed the 3D distances (annotated as *d_nei_*_gℎ*bor*_) of MERFISH-labeled neighboring genomic loci across chr21 with a genomic separation *l_nei_*_gℎ*bor*_ = 50 *kb*. The median distance for the region chr21:32.45-33.35 Mb (Supplementary Fig. S6a) was found to be 256 nm, consistent with the previous report [24]. The overall distance distribution of neighboring genomic loci across chr21 followed a log-normal distribution, with a median distance of 325 nm (Supplementary Fig. S6b, left).

Next, we calculated *d_nei_*_gℎ*bor*_ of all neighboring genomic locus pairs across 2,991 traces of chr2 that were fluorescently traced with *l_nei_*_gℎ*bor*_ = 250 *kb* (Fig 2a, green line). The distribution also exhibited a log-normal pattern, with a median distance of 462 nm and a first quartile distance *d_r_* = 299 nm (Fig. 2b, Supplementary Fig. S6b, right). For subsequent analyses, we adopted *d_r_* as an intrinsic distance reference, classifying a genomic locus pair as in spatial proximity if their measured spatial distance *d* < *d_r_*. This criterion aligns with previous studies using MERFISH data at an approximately 30 kb resolution, which employed a cutoff distance of ∼ 200 nm to define chromatin interactions [25] and 500 nm for the chr21 data with 50 kb resolution to identify genomic pairs in proximity [24].

**Fig. 2.**
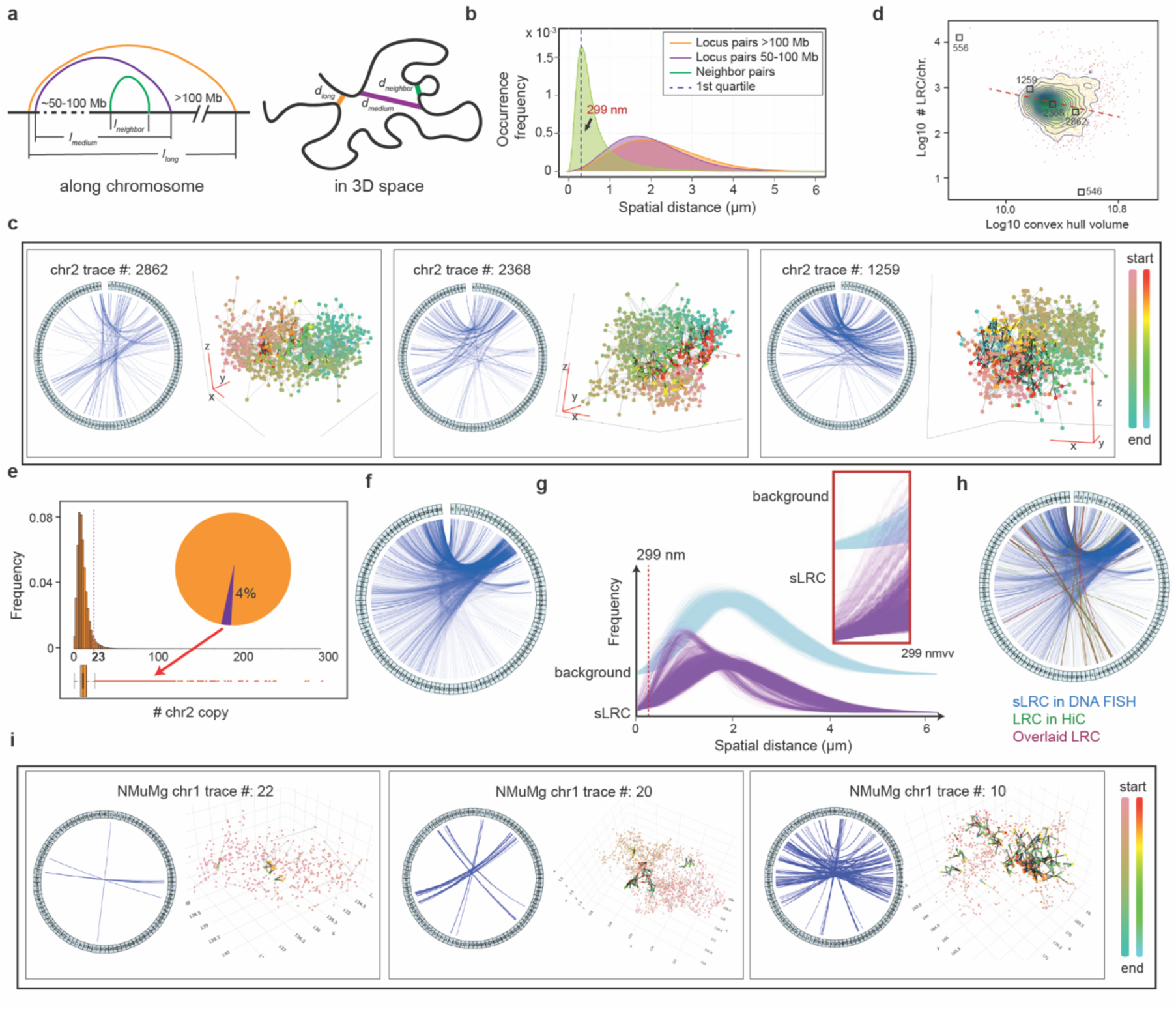
MERFISH chromatin tracing reveals widespread long-range colocalization (LRC). (**a**) Schematic illustration of genomic pairs and their genomic versus spatial distances. (**b**) Density curves of spatial distance distributions for loci pairs with genomic separations of 250 kb, 50-100 Mb, and >100 Mb. The dashed line represents the first quantile of neighboring locus pairs. (**c**) LRC linkage maps and 3D structures of 3 representative human chr 2. (**d**) Scatter plot showing the relationship between chromosome compaction, quantified by the convex hull volume, and the number of LRCs. (**e**) Histogram (top) and boxplot (bottom) showing the distribution of chr2 trace number (n = 2991) containing LRCs. LRCs present in >23 traces (red dashed line) are classified as stable LRCs (sLRCs). The pie diagram shows the percentage of LRCs classified as sLRCs. (**f**) Identification of chr2 sLRCs from DNA FISH data. (**g**) Spatial distance distributions of locus pairs with genomic separation >100 Mb. Purple lines (n = 6443): sLRCs; blue lines (n = 6443): randomly selected non-sLRCs. The red dashed line represents the first quantile of spatial distances between neighbor loci on chr2 (299 nm). (**h**) Comparison of colocalized chr2 locus pairs (>100 Mb apart) identified in IMR90 cells using DNA MERFISH (blue lines) and Hi-C (green lines) data. Overlapping links were highlighted in brown. (**i**) LRC linkage maps and 3D structures of 3 representative mouse chr 1.

To detect the presence of long-range colocalization on the chromosome, here we first calculated the spatial distances between pairs of loci with genomic separations of *l_medium_* ∈ (50, 100] Mb and *l_lon_*_g_ > 100 Mb, denoted as *d_medium_* and *d_lon_*_g_, respectively (Fig. 2a, purple and orange lines). Compared to *d_nei_*_gℎ*bor*_, the distributions of corresponding *d_medium_* and *d_lon_*_g_ shifted significantly toward larger values, with median distances of 1.86 *μm* and 2.08 *μm*, respectively (Fig. 2b). Interestingly, even at such large genomic separations, certain locus pairs exhibited spatial distances < *d_r_* (Fig. 2b, arrow). In the following analyses, we focused on the genomic locus pairs with *l_lon_*_g_ > 100 Mb and *d_long_* < *d_r_,* denoting them as long-range colocalizations (LRCs).

As expected, individual chr2 traces exhibit diverse conformations (see examples in Fig. 2c, Supplementary Fig. S6c&d, and Supplementary Movies S2-S4), consistent with previous studies [25]. Moreover, a negative correlation exists between the total number of LRCs of an individual chromosome trace and its conformation measured by convex hull volume (Fig. 2d), suggesting a potential relationship between LRCs and chromosome conformation. Notably, in these conformations, LRCs either form scattered small clusters or join into larger ones, along with varying compactness of the chromatin structures. In addition, LRC counts per chr2 trace range from fewer than 10 to over 10,000, with a peak at ∼ 600 per trace. These LRCs can be formed due to transient contacts through thermal fluctuations, or via stable structures. While transient LRCs should appear infrequently among the entire cell population, stable ones are expected to recur in subpopulations of chr2 traces. To test this hypothesis, we analyzed the occurrence frequency of 155,009 unique LRCs across 2,991 chr2 traces (Fig. 2e). We identified 7,366 LRCs with occurrence frequencies beyond the 95% population interval, and defined them as stable LRCs (sLRCs) (Fig. 2e&f). Each sLRC appeared in at least 23 chr2 traces (∼0.8% of the dataset), with some found in over 10% of samples. That is, sLRCs exhibited a significantly higher probability of occurring within 299 nm compared to other LRCs (Fig. 2g).

Some sLRC detected via MERFISH were also detected with CLOD-identified colocalized pairs (>100 Mb separation) from Hi-C (Fig. 2h). Note that Hi-C and MERFISH detect proximity differently: Hi-C captures direct contact, whereas MERFISH assesses spatial distances, using a 299 nm threshold here. Technical limitations also contribute to discrepancies — Hi-C may miss repetitive or low-accessibility regions, whereas its much larger sample size (∼1 million cells) could detect rare configurations potentially overlooked by MERFISH (with ∼3,000 chr2 traces). Despite these differences, strong agreement between Hi-C and MERFISH data indicates that stable LRCs are prevalent in chromatin.

To further validate these findings, we analyzed the locus-pair contact frequencies of IMR90 chr2 from a micro-C dataset [26] and compared them with MERFISH results. Compared to standard Hi-C, the superior coverage per genomic bin of micro-C provides sufficient signal-to-noise ratios [27] for direct analysis of raw read counts without using additional algorithms such as CLOD. When considering micro-C data with contact counts > 10 as colocalization pairs, 556 sLRCs that were identified from MERFISH data are also present in micro-C data (Supplementary Fig. S6e, left). Even when applying stringent filtering criteria (contact counts > 20), we still identified a subset of LRCs present in both micro-C and MERFISH (Supplementary Fig. S6e, right). Therefore, both sequencing-based and imaging-based approaches consistently identified sLRCs in IMR90 chr2. In addition, LRCs and sLRCs were also present in other long chromosomes of IMR90 cells (Supplementary Fig. S6f).

To assess whether LRCs exist at the single-cell level in other species, we examined a DNA seqFISH+ dataset of chr1 of mouse NMuMg cells (Fig. 2i) [28]. Applying the same analytical framework used for MERFISH, we measured spatial distance distributions for genomic loci in the DNA seqFISH+ data separated by 25 kb (which is the resolution of the dataset), 250 kb, and over 100 Mb, respectively. Those with 25 kb have the smallest median 3D distance at 253 nm. Notably, the first quartile of spatial distances for loci 250 kb apart was 281 nm in DNA seqFISH+, closely matching the 299 nm threshold from MERFISH data (Supplementary Fig. S6g). We therefore adopted a universal LRC definition across platforms: loci with genomic separation >100 Mb and spatial proximity < 299 nm. The DNA seqFISH+ dataset provided substantially lower genomic coverage per chromosome than MERFISH, with most long-range loci pairs detected in only ∼ 200 – 300 chr1 traces out of 617 or more traces (Supplementary Fig. S6h). To identify robust colocalization, we defined stable LRCs (sLRCs) as those present in at least 10, or ∼5%, of chr1 traces having the corresponding pair loci detected (Supplementary Fig. S6h&i). All these results further demonstrate that colocalization of genomically distant loci is a universal phenomenon in long chromosomes, irrespective of species origin or chromosome identity.

### Neighboring LRCs cooperate to form stable complexes through multivalent binding

DNA MERFISH data revealed the widespread presence of sLRCs (*p* ≥ 0.8%) in IMR90 chr2 traces, with some occurring in 5 – 10% of all traces (Fig. 2e). Notably, the genomic distributions of the two ends of sLRCs are not uniform but clustered along the genome (Fig. 2f). Statistical analysis confirmed that most sLRC-associated loci have their nearest neighboring sLRC genomic loci within 500 kb (Supplementary Fig. S7a). Here we defined the genomic proximity distance between two sLRC pairs *a* (loci *a_1_*, *a_2_*) and *b* (loci at *b_1_*, *b_2_*) as *l_ab_* = *max* ((|*a*_1_ − *b*_1_|, |*a*_1_ − *b*_2_|), (|*a*_2_ − *b*_1_|, |*a*_2_ − *b*_2_|)) (Fig. 3a, Supplementary Fig. S7b). Therefore, we grouped sLRCs into metaLRCs, where each sLRC had at least one other sLRC within 250 kb at both ends (Fig. 3b). This process grouped 72% of sLRCs into 736 metaLRCs, each containing 2 to over 40 sLRCs (Supplementary Fig. S7c).

**Fig. 3.**
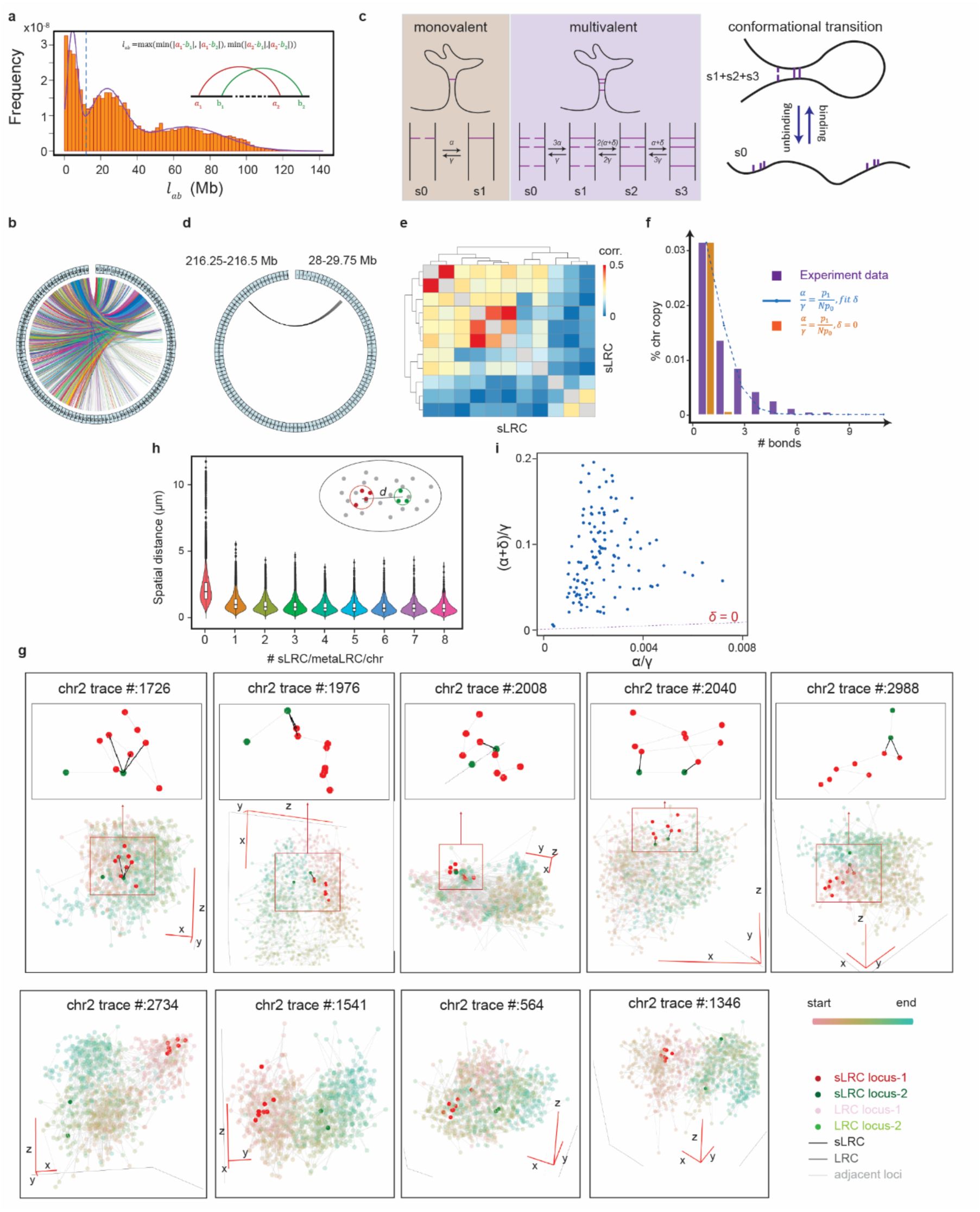
DNA FISH data analysis reveals that multivalent binding stabilizes sLRC clusters. (**a**) Frequency distribution of genomic separation between two LRCs. (**b**) sLRC and metaLRC linkage map. The outer circle represents chr2, and each line denotes a sLRC. Gray lines: isolated sLRC; colored lines: sLRCs belonging to a metaLRC. (**c**) Left: Schematic illustration of monovalent binding vs. multivalent binding models. *α*: association rate constant of a single pair in the absence of other bound sLRCs; *γ* : dissociation rate constant of a single pair; *δ* : association rate enhancement due to neighboring bound sLRCs. Right: A multi-valent site is coarse-grained into a two-state system ⎯ either free or bound (with one or more bound sLRCs) two-state system. (**d**) A representative metaLRC. (**e**) Heatmap showing correlation between sLRCs within the metaLRC from panel d. (**f**) Comparison of the probabilities of observing various numbers of bound sLRCs within the metaLRC (purple bar) vs. predictions based on an independent sLRC binding model (orange bar), and a multivalent binding model (blue dashed line). *p_0_* is not shown. (**g**) Representative 3D structures of chr2, highlighting the metaLRC from panel d. Top: Structures with at least one bound sLRC within the metaLRC. Bottom: Structures with no bound sLRC within the metaLRC. (**h**) Violin plot showing the relationship between 3D distances (between centers of minimal-sized bounding spheres of the two genomic regions forming the metaLRC and the number of bound sLRCs per individual chr2 trace. (**i**) Multivalent binding model parameters obtained through fitting DNA MERFISH data.

We hypothesized that neighboring sLRCs stabilize each other, as revealed from the live-cell imaging studies (Fig. 1i–l). To examine this hypothesis, we analyzed two classes of minimal biophysical models with and without cooperativity between sLRCs within a metaLRC (Methods). In both models, colocalization of an sLRC pair is assumed to be held by some interactions, e.g., mediated by proteins or noncoding RNAs. Consequently, a pair exists in either a bound or unbound state, distinguished by whether the two loci colocalize. In the absence of other sLRC pairs, an isolated sLRC pair stochastically transits between the two states with association (α) and dissociation (γ) rate constants (Fig. 3c, left). For simplicity, assuming that the *N* sLRC pairs within a metaLRC have the same dynamics and transit independently, one can estimate from the data 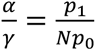, where *p_0_* and *p_1_* are the frequencies of observing zero or one bound sLRC, respectively. The probability of observing *i > 1* bound sLRC within the metaLRC can then be predicted without any adjustable parameter to be 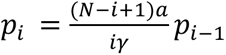.

The cooperative model assumes that the transition dynamics of neighboring sLRCs are not independent due to a multivalent binding mechanism. In the presence of other bound sLRC pairs within the metaLRC, the two loci of an unbound sLRC pair remain spatially close, leading to an increased effective association rate constant. For simplicity, assuming that the effective enhancement *δ* is the same in the presence of one or more neighboring bounded sLRC pairs, then its value can be determined by fitting the measured distribution of bound sLRCs, *p_i_* vs. *i* for i > 1. Consequently the metaLRC as a whole would appear as an effective two-conformation system with no or at least one bound sLRC, respectively (Fig. 3c, middle & right).

We tested the predictions of the two models against the MERFISH data. Figure 3d depicts one representative metaLRC. Correlation analyses over MERFISH data confirmed the cooperativity between sLRCs within the metaLRC (Fig. 3e). Note that some sLRCs have higher correlations than others within the same metaLRC, indicating that the assumption of a single value of *δ* is only crudely valid. The model without cooperativity significantly underestimated the probabilities of *p_i_* for i > 1, while a single parameter *δ* > 0 significantly improves the agreement between the experimental data and theoretical predictions. Two additional examples shown in the Supplementary Fig. S7d&e also support the cooperative model. The bimodal distribution in Supplementary Fig. S7d reveals an even stronger cooperativity than the simple model we considered, suggesting that the enhancement *δ* increases with the number of bounded sLRCs.

Analysis of individual chr2 trace further supported the cooperative model. The spatial distances between two tagged genomically distant regions in bound form were much shorter compared to those in free form chr2 traces (Fig. 3g, Supplementary Movies S5–8). When one or more sLRCs were bound, the two genomic regions within the metaLRC were held closer, while additional bound sLRCs (> 1) did not significantly alter spatial distance (Fig. 3h). The observed non-monotonic *p_i_* vs. *i* curves for certain representative metaLRCs (e.g., 33.75–35.5 Mb & 146–146.25 Mb; 119.25–123.75 Mb & 242 Mb) suggested even stronger cooperativity than the minimal model predicted (see Methods) (Supplementary Fig. S7d&e).

Next, we examined all metaLRCs of chr2 with *N* > 2. The *α*/*γ* values directly obtained from the FISH data were < 0.007 (Fig. 3i), indicating that associating two distant loci is entropically unfavorable. However, the corresponding (*α* + *δ*)/*γ* values were up to 10-100× higher, making the probability of observing a bound metaLRC over 1-10% (Supplementary Fig. S7f). That is, pairs of distant loci can remain stably proximate within a chromosome via neighbor-assisted multivalent binding, significantly enhancing structural stability and long-range genomic interactions.

### Structural analyses reveal multiple candidate mechanisms for sLRC formation

Colocalization of distant cis-loci has been intermittently observed. Three general mechanisms have been proposed to explain LRC formation (Fig. 4a, I–III): (I) loose/compact region segregation, may bring distant compact region loci into transient proximity [29], or relatively stable colocalization via polycomb structures [12]; (II) Transcriptionally active loci, around which chromatin is typically loose and accessible, may colocalize via attaching to shared nuclear structures like speckles or nucleoli [30]; (III) The nuclear membrane may recruit multiple loci into spatial proximity [31]. In addition, one may conceive mechanisms in which sLRC formation is stable and independent of major nuclear structures or compartment segregation. Here, we generalize them as Mechanisms IV. While the first three have been proposed previously [32], sLRC assemblies formed through Mechanism IV were considered rare.

**Fig. 4.**
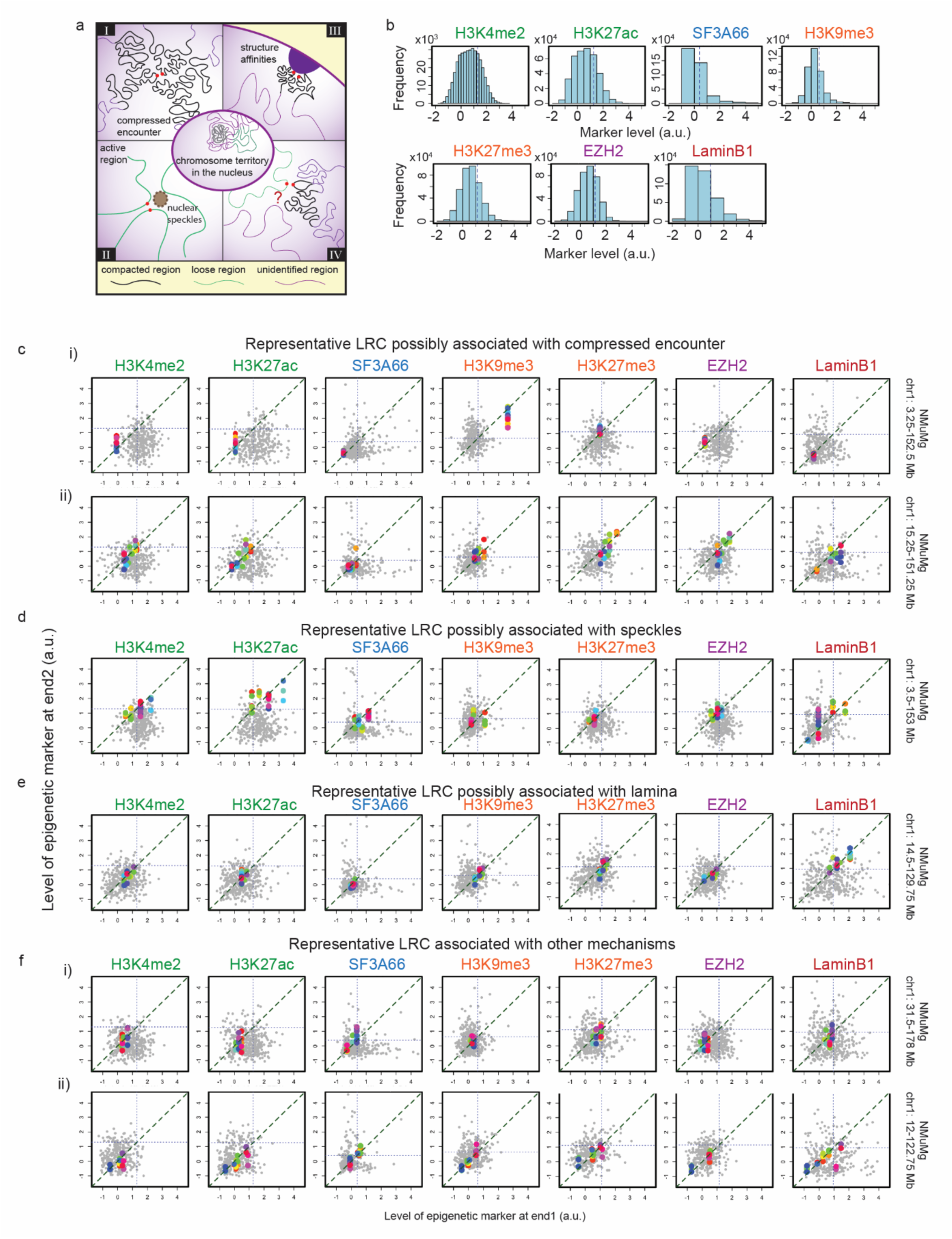
Possible mechanisms of sLRC formation and example LRCs from the DNA seqFISH+ data with epigenetic mark patterns consistent with different mechanisms. (**a**) Schematic of four possible mechanisms for sLRC formation. I and II: loose/compact region segregation leads to compressed encounters of genomically distant loci in the compact and loose regions, respectively. III: Colocalization of transcriptionally inactive, genomically distant loci interacting with common nuclear membrane structures. IV: A subset of sLRCs shows no epigenomic features consistent with Mechanisms I–III, suggesting the involvement of currently uncharacterized interactions. This category is defined by exclusion and its molecular basis remains an open question. (**b**) The histograms depict the distributions of epigenetic levels on all detected loci. (**c-f**) Example LRCs with diverse epigenetic patterns. Every point on the plot depicts one pair of loci on a single chromosome. The grey points are those chromosomes without colocalization, and the color points are those with colocalization.

To explore possible mechanisms of sLRC formation, we first investigated bulk A/B compartment annotations for IMR90 cells [15], with sRLCs identified by MERFISH (details in Methods). The analysis revealed no clear pattern or correlation between the two loci of each sLRC pair (Fig. S8a, left), whereas a large subset of sLRCs have both ends in the chromatin-loose region (Fig. S8a, right), consistent with a previous study [24]. To further validate the results, we also investigated another CHARM dataset [33] (details in the Methods). Similarly, most LRCs show no clear pattern (Fig. S8b). Together, our integrative analysis across IMR90 and mESC datasets did not reveal any clear preference of LRC formation for specific A/B compartment combinations. This once again highlights the complexity of LRC formation, suggesting that mechanisms beyond known regulators are involved. However, such conclusion should be taken with caution since the A/B compartment was calculated from bulk/pseudo-bulk-level data. That is, an ensemble of chromosomes may be composed of a mixture of subpopulations with and without the presence of an sLRC under study.

Therefore, we linked the sLRC and associated loci to epigenetic marks and nuclear structures at the single-chromosome level using the DNA seqFISH+ dataset [28]. This dataset with single-chromosome resolution combines spatial coordinate information of each location on the mouse genome with its epigenetic modification marks, such as H3K4me2, H3K27ac, H3K9me3, H3K27me3, etc., as well as marks for nuclear structures, such as those for Polycomb (EZH2), speckles (SF3A66), and lamina-associated domains (Lamin B1). To be consistent with the resolution of MERFISH, we grouped the DNA seqFISH+ data to a 250 kb resolution, calculated the median mark level, and used the 3^rd^ quantile as the gate to identify whether a region has a high level of the targeted marker (Fig. 4b).

The epigenetic profiling reveals a subset (43%) of sLRCs exhibiting epigenomic features consistent with the three identified formation mechanisms (Mechanisms I–III, Fig. 4a). For instance, among the 323 detected traces of the 3.25 Mb and 125.15 Mb locus pair on NMuMg chr1, 15 colocalized traces exhibited elevated H3K9me3 and uniformly depleted euchromatin marks (H3K4me2 and H3K27ac) relative to non-colocalized traces (Fig. 4c-i). Another pair (15.25 Mb and 151.25 Mb) displayed high H3K27me3 enrichment along with the Polycomb marker EZH2 (Fig. 4c–ii). Both examples are consistent with compressed encounters driven by A/B compartment segregation (Mechanism I). A colocalized pair in Fig. 4d (3.5 Mb and 153 Mb) instead showed high levels of active chromatin markers (H3K4me2 and H3K27ac), enrichment of the nuclear speckle marker SF3A66, and low H3K9me3 and H3K27me3, consistent with nuclear speckle–associated assembly (Mechanism II). Fig. 4e illustrates an example potentially following Mechanism III, with high Lamin B1 enrichment indicating association with lamina-associated domains.

Notably, over half of sLRCs (57%) on mouse chr1 could not be assigned to these categories: the regional epigenetic mark levels of the two ends of an sLRC show no strong correlation (Fig. 4f–i); or all epigenetic mark levels of both ends are relatively low or show no clear pattern (Fig. 4f–ii). These observations are inconsistent with Mechanisms I–III, suggesting that at least a subset of sLRCs may arise through one or more additional pathways not captured by the current epigenetic markers examined. We provisionally label this open category Mechanism IV, while acknowledging that the molecular basis of such interactions remains unknown and may be heterogeneous. Definitive characterization will require perturbation experiments targeting candidate factors such as bridging proteins, repetitive elements, or non-coding RNAs.

While features of individual loci can be identified from the key epigenetic marks of the DNA FISH+ data, there is no epigenetic information at the single-chromosome level for the MERFISH data. As an alternative, we examined the local density features of a locus, noting that the two ends of an sLRC formed through Mechanism I-III are expected to share similar local packing structures. Specifically, by defining regional density of short-range neighbors (RD-SN) and regional density heterogeneity (RDH), which measures the uniformity of local spatial distributions (Supplementary Fig. S9a-c and Methods). Then one expects that Mechanism I and III lead to high RD-SN for both loci associated with an sLRC, Mechanism II leads to low-low RD-SN, while the pattern for Mechanism IV is undetermined. Furthermore, a locus next to a nuclear structure (Mechanism II and III) is expected to exhibit high RDH due to the excluded volume of the latter. Therefore, a combined RD-SN and RDH analysis can identify sLRCs not convincingly explainable by Mechanisms I-III, e.g., those with both RD-SN and RDH low on both loci. Indeed, sLRCs show combinatorial patterns of these two quantities, implying the existence of diverse mechanisms for sLRC formation (Supplementary Fig. S9d&e), as also reflected by examining representative structures (Supplementary Fig. S9f).

Furthermore, we assessed the relationship between LRCs and cell cycle phases. To do this, we investigated the CHARM dataset that simultaneously profiled 3D chromosome structure and the transcriptome at single-allele resolution [33]. We first identified pairs of long-range loci separated by more than 100 Mb, retaining only those pairs that established physical contact in more than 20 out of 959 cells for subsequent statistical analysis. Then, we inferred the cell cycle phase of every single cell directly from the transcriptomic data based on the expression of canonical S-phase and G2/M-phase marker genes. For each LRC, contact enrichment for each phase was calculated as the proportion of cells in that phase that harbor the contact. Significant pairs were defined as those with P < 0.05. Of the 149 valid LRCs (Fig. S8c), we found only 11 pairs with significant enrichment or depletion of contacts in specific cell-cycle phases (P < 0.05) with distinct patterns (Fig. S8d). Among them, nine LRCs were preferentially established during G2/M, the other two during G1 phase. All these findings suggest that most long-range contacts are also independent of the cell cycle stage.

In addition, we also investigated possible functions of sLRC in gene expression. Use the same CHARM dataset, we examined the genes located near two endpoints of the targeted locus pairs and classified the alleles into two groups: contact alleles (those in which the two ends of the locus pair form a LRC) and no-contact alleles (those in which they are not). scHiC-defined allele pairs with and without contacts show a strong correlation in mean expression for most genes (Fig. S8e), indicating that LRC formation between most long-range locus pairs does not significantly affect gene expression. To further investigate this relationship, we examined a selected subset of locus pairs in detail. In most pairs of long-range loci, genes on the loci showed no clear pattern of change in expression level with or without contact (Fig. S8f). Some pairs did show distinct expression patterns with and without contact. Upon contact formation, some genes anchored at certain pair showed decreased expression compared to the non-contact case (Fig. S8g). Such behavior is consistent with compressed encounter (Mechanism I). Conversely, for some pairs of loci, when contacts were established, genes on them showed elevated expression compared to those on no-contact alleles (Fig. S8h). Such observations are consistent with Mechanism II, e.g., the contacting loci are recruited to a common nuclear speckle. Together, these analyses provide evidence that formation of only some LRCs correlates with gene co-expression and transcriptional regulation.

Collectively, all structural and functional results point to multiple mechanisms and possible functions underlying sLRC stability. While three of these mechanisms have been well characterized, a previously underappreciated one emerges as an additional contributor to stable sLRC formation.

## Discussion

Our analyses of Hi-C and DNA FISH chromosome tracing data reveal widespread colocalization of genomic loci separated by distances far exceeding typical TAD distances. It is important to note that our study employed stringent criteria to identify representative examples rather than exhaustively cataloging all stable long-range colocalizations.

Statistically, the probability of multiple genomic loci with *l* ≥ 100 Mb being in physical proximity without stable structural mechanisms would be negligible. Some sLRCs may form through known mechanisms such as compartment segregation or anchoring to nuclear structures such as nucleoli, speckles, or the nuclear envelope. However, many sLRCs are buried in the nuclear interior, show no signature of compartment segregation, and are away from large nuclear structures. While this pattern is inconsistent with Mechanisms I–III, it is important to note that the absence of known signatures does not by itself constitute evidence for a distinct mechanism. We therefore use ’Mechanism IV’ as a placeholder category for sLRCs that current epigenomic and spatial data cannot explain, rather than as a validated mechanistic class. Key open questions include: what molecular interactions hold these assemblies together; whether the category is mechanistically unified or heterogeneous; and whether candidate elements such as repetitive sequences, non-coding RNAs like LINC01317, or multivalent bridging proteins are causally involved. Future perturbation studies, such as targeted depletion of candidate factors or deletion of repetitive elements at hub loci, will be necessary to establish causality.

The molecular nature of the interactions holding the assemblies needs further investigation. An assembly can be liquid-, gel-, or solid-like, similar to what is observed in other contexts. Interestingly, there are some loci that function as integrative hubs, colocalizing with multiple genomically distinct loci with various epigenetic states. This suggests that they may mediate the assembly of LRCs differently than their partnering loci. These hub sites can either facilitate liquid droplet formation or act as scaffolds for solid assembly mediated by elements such as non-coding RNAs or bridging proteins. For instance, the human chr2 242 Mb locus is enriched in repetitive sequences, which may serve as selective binding sites for partnering loci, either directly or through other bridging factors. Another 32–35 Mb locus on human chr2 is also rich in repetitive sequences and non-coding RNAs such as LINC01317. Multi-way chromosome assemblies have been identified in different contexts, such as enhancer-promoter hubs [34], superenhancers, and the multi-chromosome enhancer clusters controlling monoallelic olfactory receptor expression in olfactory sensory neurons [12, 28, 35, 36]. The assemblies of sLRCs and metaLRCs may form under similar molecular mechanisms. One known example is that Polycomb repressive complex 2 uses RNA binding to coordinate long-range chromosome loop formation and H3K27me3 spreading [12]. Falk et al. argue that attractions between heterochromatic regions and lamina-chromatin interactions, i.e., Mechanisms I and III, are critical to recapitulate compartmented chromatin structures [37]. Analyses in this work reveal existence of long-range colocalization structures showing no clear features of lamina-chromatin interactions and heterochromatin, and their molecular mechanism(s) of formation need further investigation.

A key unsolved question is how sLRC loci locate and associate with each other. One plausible mechanism is that, during post-mitotic DNA decondensation, some genomically distant loci may be in spatial proximity and associate stochastically, leading to subsequent formation of additional sLRCs and stabilization of the assembled structures. Liquid-phase droplet fusion can also drive two regions into proximity to form an sLRC [38].

Besides their varied formation mechanisms, we hypothesize that these sLRC structures play diverse functional roles, including transcriptional regulation and broader chromosome 3D organization. Proper DNA folding into spatially segregated territories with minimal entanglement and knot formation is essential for eukaryotic genome organization and gene regulation. This process must be robust across all cell types, including those with abnormal karyotypes, and must function despite cell type-specific euchromatin/heterochromatin partitioning, folding stochasticity, and mutations. From an engineering perspective, achieving this through a single centralized mechanism requiring precise tuning would be challenging. Instead, our findings suggest a redundant, distributed component mechanism that facilitates chromosome folding into compact structures (Fig. 5). The genome harbors an extensive repertoire of NCs and partnering loci separated by long genomic distances, from which only a subset is available for a given cell type. At the individual chromosome level, subsets of these loci stochastically form sLRCs and larger structures, contributing to the observed heterogeneity of chromosome structures [39]. These structures provide a “divide-and-conquer” mechanism to facilitate forming chromosome structures at smaller length scales, e.g., by forming segregated globally compact chromosome configurations, and closed-end boundary conditions for simultaneous crumpled folding at multiple regions [5], and enhanced contact frequencies for forming loops at smaller scales.

**Fig. 5.**
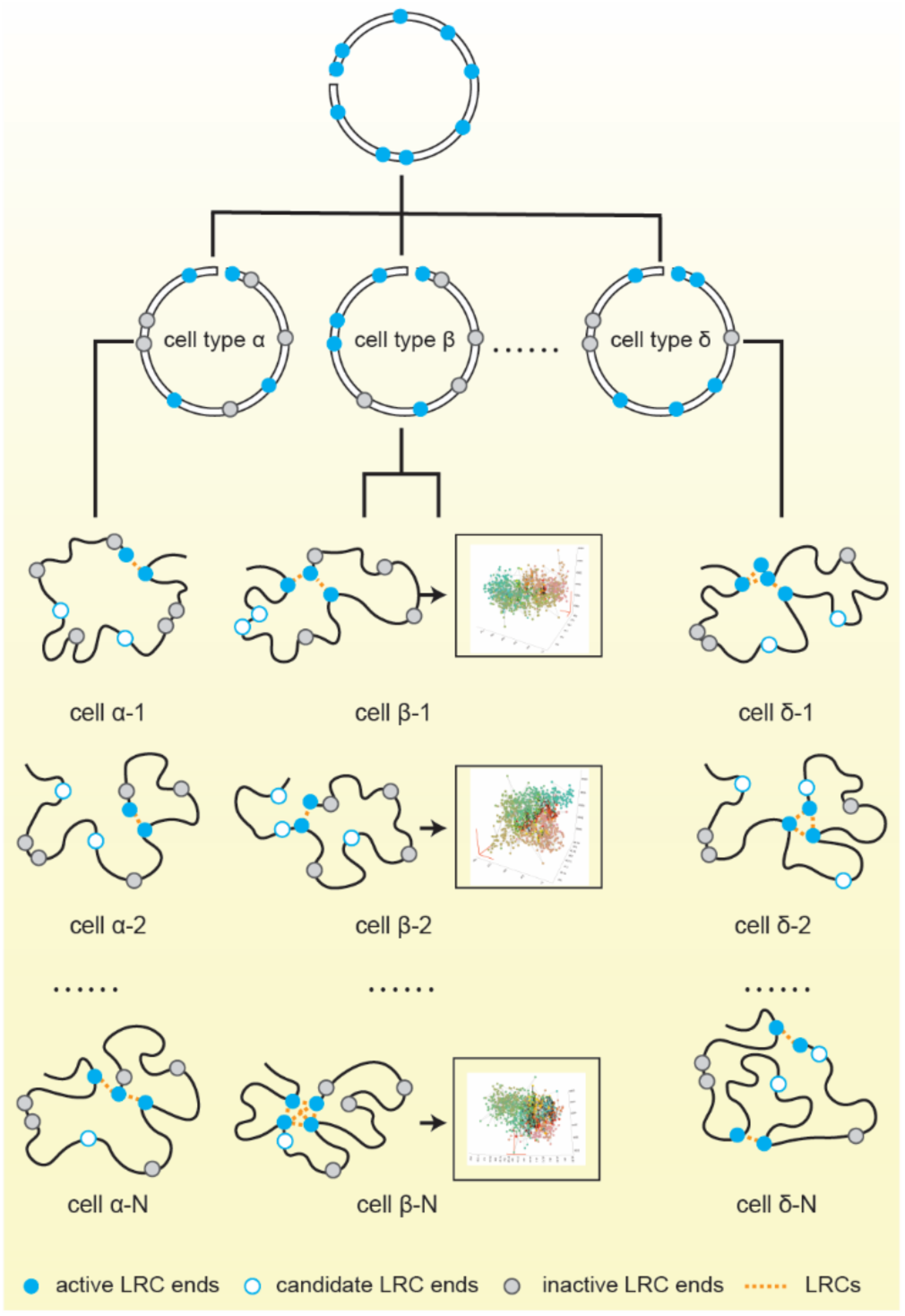
Schematic illustration of the proposed excessively redundant distributed component mechanism for chromosome folding.

## Conclusions

In summary, our analyses of chromosome tracing and Hi-C data uncover prevalent, conserved large-sized loops and their clustering into stable structures. These findings raise important questions regarding the molecular mechanisms, temporal dynamics, and functional significance of these structures on chromosome configurations, gene transcriptional activity, and cell type regulation. Importantly, while three formation mechanisms can be linked to known nuclear biology, a fourth class of stable colocalizations — identified here by the absence of known structural signatures — points to molecular interactions that remain to be discovered, and should be a focus of future experimental and computational work.

## Methods

### Hi-C data analysis

#### Raw data analysis

Raw Hi-C data of human were downloaded from GEO, as shown in Supplementary Table 1, and were aligned to the human genome *hg38* using the Burrows-Wheeler Alignment tool [40]. Hi-C contacts were then identified and filtered with Pairtools [41]. The contact matrices of balanced contact intensity used for the CLOD algorithm were created with Cooler [42]. Hi-C counts of other species were downloaded from GEO directly (Supplementary Table 1).

#### Chromosome Long-range colocalization identifier through Outlier Detection (CLOD) pipeline

To establish a minimalistic and robust pipeline to detect long-range colocalization between cis-loci based on Hi-C results, we smoothed the noisy Hi-C data and then highlighted the robust off-diagonal signals representing chromatin contacts between distal genomic regions. First, we used the balanced Hi-C results and converted the original reads to the average of the window to smooth the random, noisy signals, and to distinguish colocalization signals from background noise. In detail, we grouped all pairs of loci with the same genomic distances (*l*) and those with *l* ± *resolution* and calculated the quantiles. The final output of the CLOD is the fold change of the interquartile range above the third quantile of all pairs with similar genomic distances 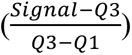 (Supplementary Fig. S3). In this paper, we defined locus pairs as those whose signals exceed two folds of the interquartile range above the third quartile of all pairs with similar genomic distances, *i.e*., 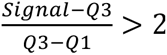.

### MERFISH data

The Euclidean coordinates of chromosome fragments were directly downloaded from Zenodo repository (records:3928890).

### DNA seqFISH+ data

The Euclidean coordinates of chromosome fragments and the level of epigenetic marks were directly downloaded from Zenodo repository (records:7693825).

To address the analytical challenges posed by the lower genomic coverage of DNA seqFISH+ data, we developed a conservative aggregation strategy that also ensures methodological consistency with the 250 kb resolution used in MERFISH analysis. Linear interpolation was considered but avoided, as it may introduce artificial dependencies between loci. Instead, we divided the genome into consecutive 250 kb bins, with each bin boundary serving as a genomic node. For each such node, we aggregated all original detected loci located within a 100 kb flanking window (i.e., ±100 kb around the node). Two nodes were defined as being in contact if any pair of original loci between the two coarse-grained regions fell within the established proximity threshold. This aggregation strategy enhances the effective coverage of the DNA seqFISH+ data and reduces sparsity while maintaining the fidelity of the underlying biological signal.

### CRISPR-dCas9 two-color labeling

The plasmid cloning method used for CRISPR-dCas9 fluorescent labeling was adapted from a previously established protocol [43]. The sgRNAs used in this study are listed in Supplementary Table 2.

#### Cell culture

HEK293T cells (#ATCC-CRL-3216) were cultured in DMEM (Invitrogen #11995073) supplemented with 10% FBS (Gbico #10437028), 100 units/ml penicillin, and 100 µg/ml streptomycin (Invitrogen). Cells were maintained at 37°C in 5% CO2, and used within 15–30 passages for experiments.

For transfection, HEK293T cells were seeded onto 35 mm glass-bottom dishes one day before the experiments. In the next day, 750 ng of donor template and 750 ng of sgRNA-Cas9 plasmid were co-transfected using FuGENE HD Transfection Reagent (Promega #E2311). Cells were incubated for 48 hours before imaging[43].

#### Time-lapse imaging

For imaging presentation, cells were visualized using a Nikon Ti microscope equipped with a 100x oil objective. Excitation wavelengths were 470 nm for GFP and 555 nm for RFP. Cells were imaged every 10 seconds for 10 minutes with an exposure time of 50 ms.

For trajectory tracking, cells were maintained in a Tokai Hit Microscope Stage Top Incubator and imaged using a Nikon Ti microscope with FITC and TRITC channels (excitation wavelength is 470 nm and 555 nm for GFP and RFP, respectively; 60x oil objective). Cells were imaged every minute for 60 minutes with an exposure time of 100 ms [44].

#### Imaging processing

Due to the low number of cells, cell segmentation and puncta recognition were performed by CellProfiler [45], and mislabeled points were corrected manually.

### Derivation of the minimal multivalent binding model

Consider a metaLRC that contains *N* pairs of sLRCs. Given the observed abnormally high contact frequency of a sLRC pair as compared to non-sLRC pairs separated with similar genomic distance, it is likely that the two loci are held together through some direct interactions or mediated by other molecular components. Thus, we can assume that an sLRC pair *i* exists in one of the two possible configurations, *σ_i_* = 1 if the pair is bound (with spatial distance *d_r_* < = 299 nm), and 0 otherwise. We assume this metaLRC is genomically distant from other metaLRCs and sLRCs, so for a good approximation there is no need to explicitly consider the influence of the latter on the association-dissociation dynamics of sLRCs in this tagged metaLRC.

Suppose that without influence from other sLRCs within the tagged metaLRC, an sLRC pair has a bare association rate *α* and a dissociation rate γ. Then at kinetic equilibrium, the probability in the bound state is 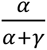, and from the FISH data we expect that *α* ≪ *γ*. Here for simplicity we assume that each sLRC pair has the same association-dissociation kinetic parameters. Let’s focus on one pair *i*. When there is one or more other bound sLRC pairs within the metaLRC, after dissociation the two loci of pair *i* are constrained by the nearby bound sLRCs and cannot diffuse far away from each other before rebind. Note that the genomic distance between these two loci of a sLRC ≥ 100 Mb, and by definition the genomic distance between two sLRCs *i* and *j* within a metaLRC *l_ij_* << 100 Mb. Consequently, at the presence of other bound sLRC pair(s), the two loci of a dissociated sLRC have an increased probability to confront each other and rebind, so the effective association constant changes to *α* + *δ*, and we expect that *δ* ≫ *α*. For simplicity, given *l_ij_* << 100 Mb let us assume that each sLRC pair has the same effect on any other one, and the effect is not additive but *δ* remains the same value at the presence of one or more bound sLRC pairs. Given the genomic distance between two loci belonging to two different sLRCs, we further assume that there is no direct interaction between two sLRC pairs to simplify the analyses. That is, we assume that the dissociation constant of a sLRC pair, γ, is not affected by the presence of other bound sLRC pairs.

With the above model, one can write down a set of master equation for finding that the metaLRC exists with 0, 1,…, *N* pairs of bound sLRCs,

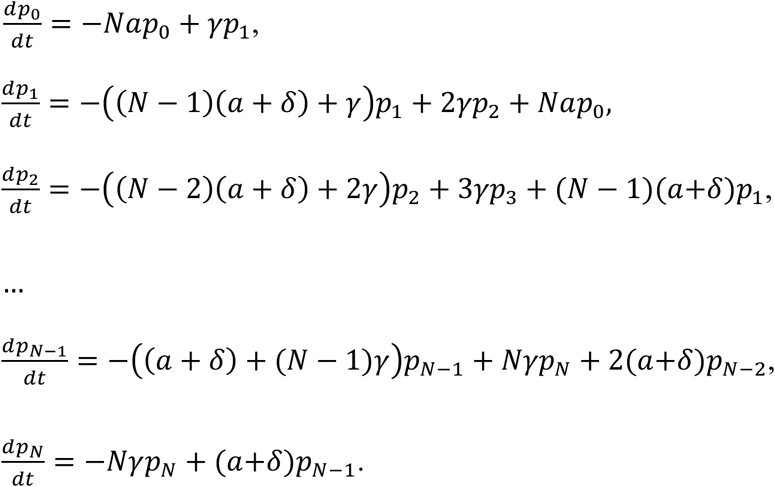

Then at steady state,

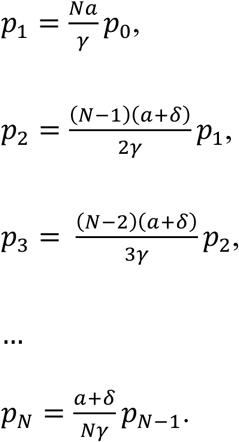

Define the probability of observing presence of one or more pairs as

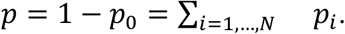

One has,

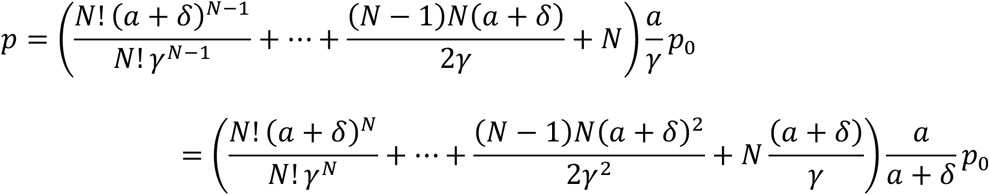

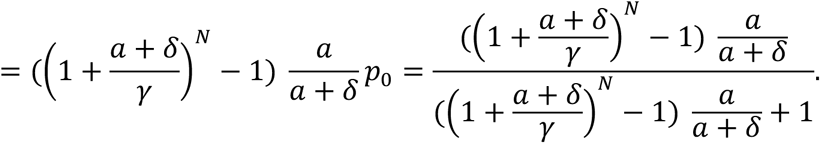

In the limit *α* ≪ *δ* ≪ *γ*, the above expression can be further simplified as,

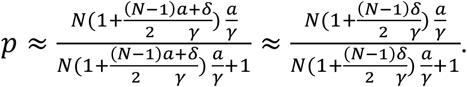

Several mechanisms can further increase the cooperativity beyond what described by a constant *δ* value in the above model. Presence of additional bound sLRC pair may further constrain the diffusion of the two loci of a dissociated sLRC, leading to further increase of the value of *δ*. That is, the value of *δ* may increase with the number of bound sLRC pairs. The bound sLRC pairs may physically contact and stabilize each other with an increased *γ* value. Indeed, we observed non-monotonic histograms of *p_i_* for some metaLRCs (Supplementary Fig. S7d&e). With a thermodynamic equilibrium model, one can introduce some binding free energies to describe the cooperativity as in allosteric effect and a previous cooperative enhancer binding model [35]. For simplicity in this work, we restricted the analyses to the minimal model discussed above.

Note that all the loci belonging to a metaLRC can be clustered as two clumped genomic regions that are genomically distant (i.e., ∼ 100 Mb or further). Subsequent analyses further coarse-grained the binding state of a metaLRC as 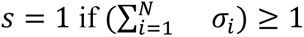 and 0 otherwise.

### A/B compartment assignment

For the bulk A/B compartment annotations from IMR90 cells [15], bins with PC1 ≥ 0.3, PC1 ≤ −0.3, and |PC1| < 0.3 were classified as A, B, and boundary, respectively.

For the pseudo-bulk level A/B compartment from CHARM data set [33], pseudo-bulk Hi-C contact matrices were generated for chr 1, normalized by observed/expected (O/E), and subjected to PCA to derive PC1 and normalized. Bins were classified as A (PC1 > 0.3), B (PC1 < −0.3), or boundary (|PC1| ≤ 0.3).

### Quantification of local geometric structures from MERFISH data

The local geometric structures of every locus from MERFISH data were identified by the <u>R</u>egional <u>D</u>ensity of <u>S</u>hort-range Neighbors (RD-SN) and the <u>R</u>egional <u>D</u>ensity <u>H</u>eterogeneity (RDH). In detail, in the MERFISH data, following the 299 nm threshold used in this paper, we define the sphere space with a radius of 299 nm as the region around a locus (Supplementary Fig. S9a). For RD-SN, we calculated the number of other loci that are within the region and less than 10 Mb from the center locus (Supplementary Fig. S9b, top). For RDH, we first equally divide the region sphere into eight sections and then calculate the variance of the number of loci in each section (Supplementary Fig. S9c, top).

### LRC identification from the CHARM data

We investigated allele-specific scHiC maps at 250 kb resolution in conjunction with transcriptomic expression data. We first identified pairs of long-range loci separated by more than 100 Mb, retaining only those pairs that established physical contact in more than 20 out of 959 cells for subsequent statistical analysis.

### Gene expression analyses from the CHARM data

For each retained locus pair mentioned before, we examined the genes located near its two endpoints, and classified the alleles into two groups: contact alleles (those in which the two ends of the locus pair are contacted) no-contact alleles. For every gene, we then computed the mean expression level separately for contact alleles and no-contact alleles,

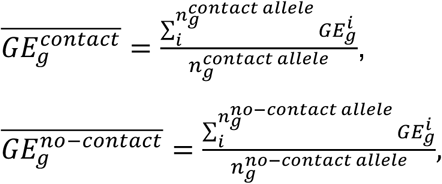

where 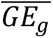 is the mean expression of gene *g* in contact or no-contact alleles, 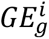 the expression level of gene *g* on allele *i*, and 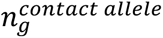 and 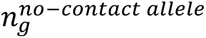 are the total number of observed alleles of gene *g* with and without LRC formation, respectively.

## Abbreviations

LRC: Long-range colocalization for genomic locus pairs > 100 Mb and within 299 nm in spatial distance.
sLRC: Stable long-range colocalization for LRC pairs with occurrence frequency > 4% in the IMR90 chr2 MERFISH dataset, or the top 5% in IMR90 chr1, chr3 and chr4 MERFISH dataset, or ≥15 (∼10% in detected chromosomes) in NmuMg chr1 DNA seqFISH+.
metaLRC: a group of sLRCs, in which for each sLRC there was at least one other sLRC with 250 kb for both ends.
NC: Nucleation center referring to locus having multiple sLRC partner loci.
RD-SN: Regional density of short-ranged neighbors (< 10 Mb) within a 299 nm range of a tagged locus.
RDH: Regional density heterogeneity.

## Authors’ contributions

Conceptualization: JZ, JX

Methodology: JZ, SW, SCW, JX

Investigation: JZ, JX

Visualization: JZ, JX

Funding acquisition: JX

Project administration: JX

Supervision: JX

Writing – original draft: JX

Writing – review & editing: JZ, SW, SCW, JX

## Acknowledgments

We thank Tom Mistelli, Harinder Singh, Jing Chen, Jian Liu for helpful discussions.

## Code availability

The codes used in this paper were submitted to https://github.com/xing-lab-pitt/LRC-notebooks.

## Data availability

All data used in this paper are listed in Table S1.

## Supplementary Materials

**Supplementary Table S1.** HiC datasets used in CLOD analyses.

| Cell type | Dataset source | References |
| --- | --- | --- |
| ESC | GSE52457 | 18 |
| ME | GSE52457 | 18 |
| MSC | GSE52457 | 18 |
| NPC | GSE52457 | 18 |
| RWRE1 | GSE118629 | 19 |
| C42B | GSE118629 | 19 |
| HPNE | GSE149103 | 20 |
| MCF7 | GSE66733 | 14 |
| MCF10A | GSE66733 | 14 |
| 22Rv1 | GSE118629 | 19 |
| Capan-1 | GSE149103 | 20 |
| PANC-1 | GSE149103 | 20 |
| IMR90 | GSE63525 | 15 |
| HEK293T | GSE44267 | 23 |
| HCT116 | GSE104333 | 17 |
| Mouse-lung | GSE295452 | 21 |
| Mouse embryo | GSE167579 | 22 |
| Rat | GSE167579 | 22 |
| Rhesus | GSE167579 | 22 |
| Cattle | GSE167579 | 22 |
| Sheep | GSE167579 | 22 |
| Rabbit | GSE167579 | 22 |
| Chicken | GSE167579 | 22 |

**Supplementary Table 2.**
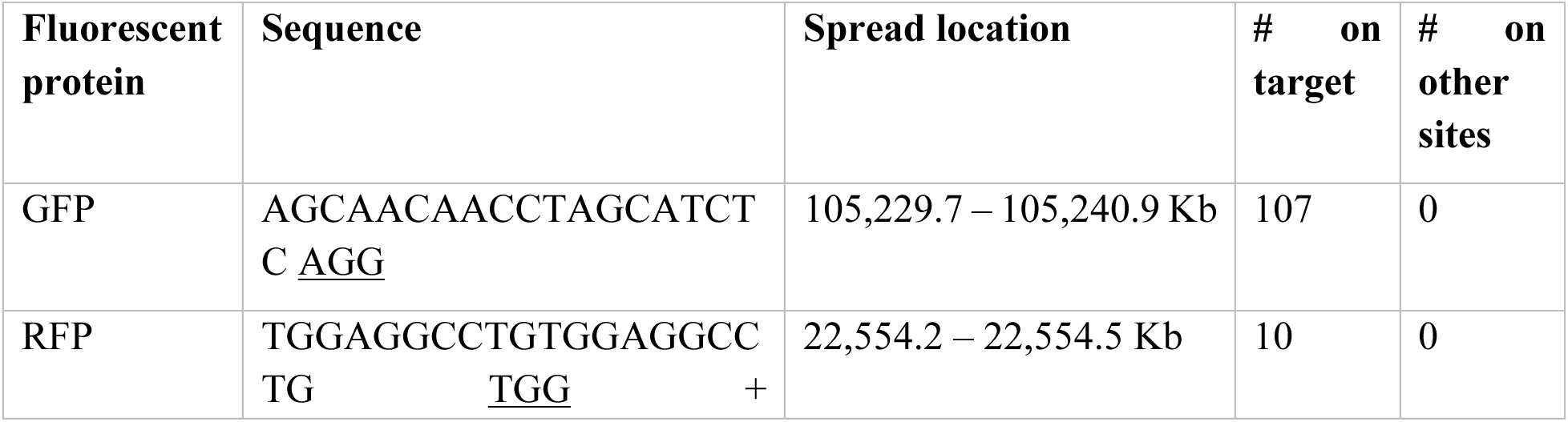
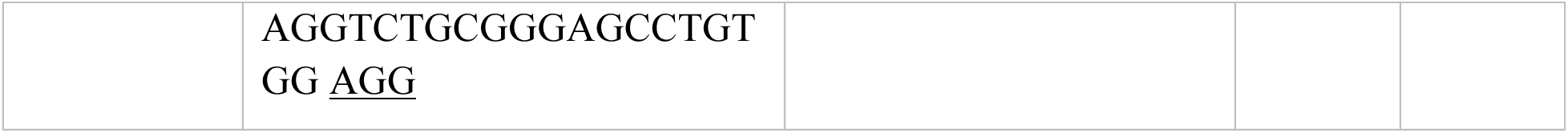
Summary of tested sgRNA.

**Supplementary Movie S1.**

HEK293T CRISPR knock-in fluorescent microscopy imaging.

**Supplementary Movies S2-S4**

3D structure of representative chr2 traces.

**Supplementary Movies S5-S8**

Comparison of 3D structure of example chr2 traces with and without metaLRC.

**Supplementary Fig. S1.**
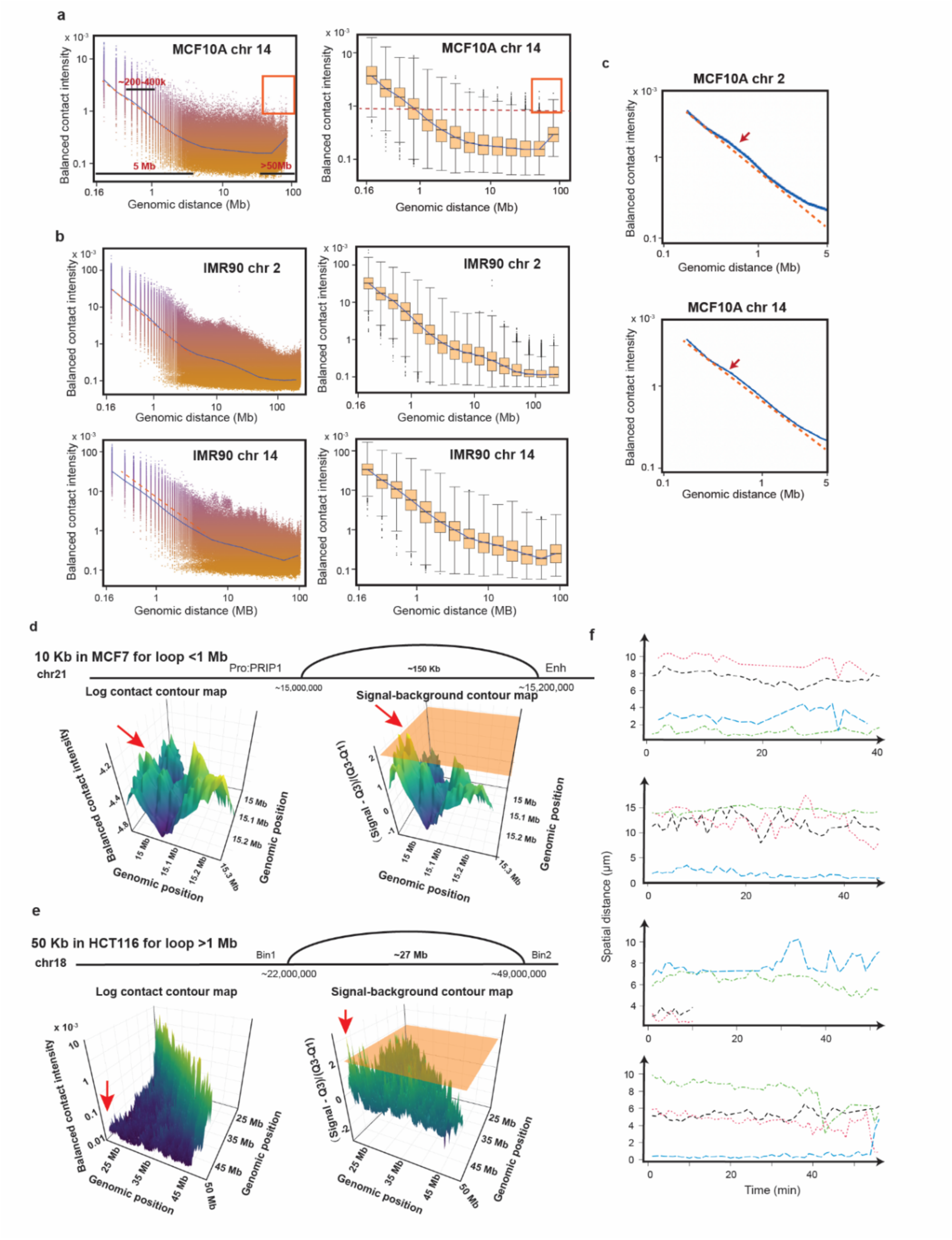
Decay trends of contact frequency in additional cell lines and CLOD pipeline. (**a**) Scatter plot (left) and boxplot (right) showing the relationship between contact frequency and genomic distance for chr14 in MCF10A cells (blue line). Red-boxed points represent locus pairs separated by > 50 Mb, exhibiting contact frequencies higher than the third quartile of pairs separated by ∼1 Mb. (**b**) Same as panel (a) but for chr2 and chr14 of IMR90 cells. (**c**) Zoomed-in view of the contact frequency vs. genomic distance trend, derived from Fig. 1a and panel a. (**d**) Experimental validation of colocalized pairs identified by CLOD, using MCF7 cell Hi-C data at 10 kb resolution. (**e**) Same as panel d but using HCT116 cell Hi-C data at 50 kb resolution. (**f**) The distances between blue and red puncta pairs in each of the four cells that were tracked for 60 min.

**Supplementary Fig. S2.**
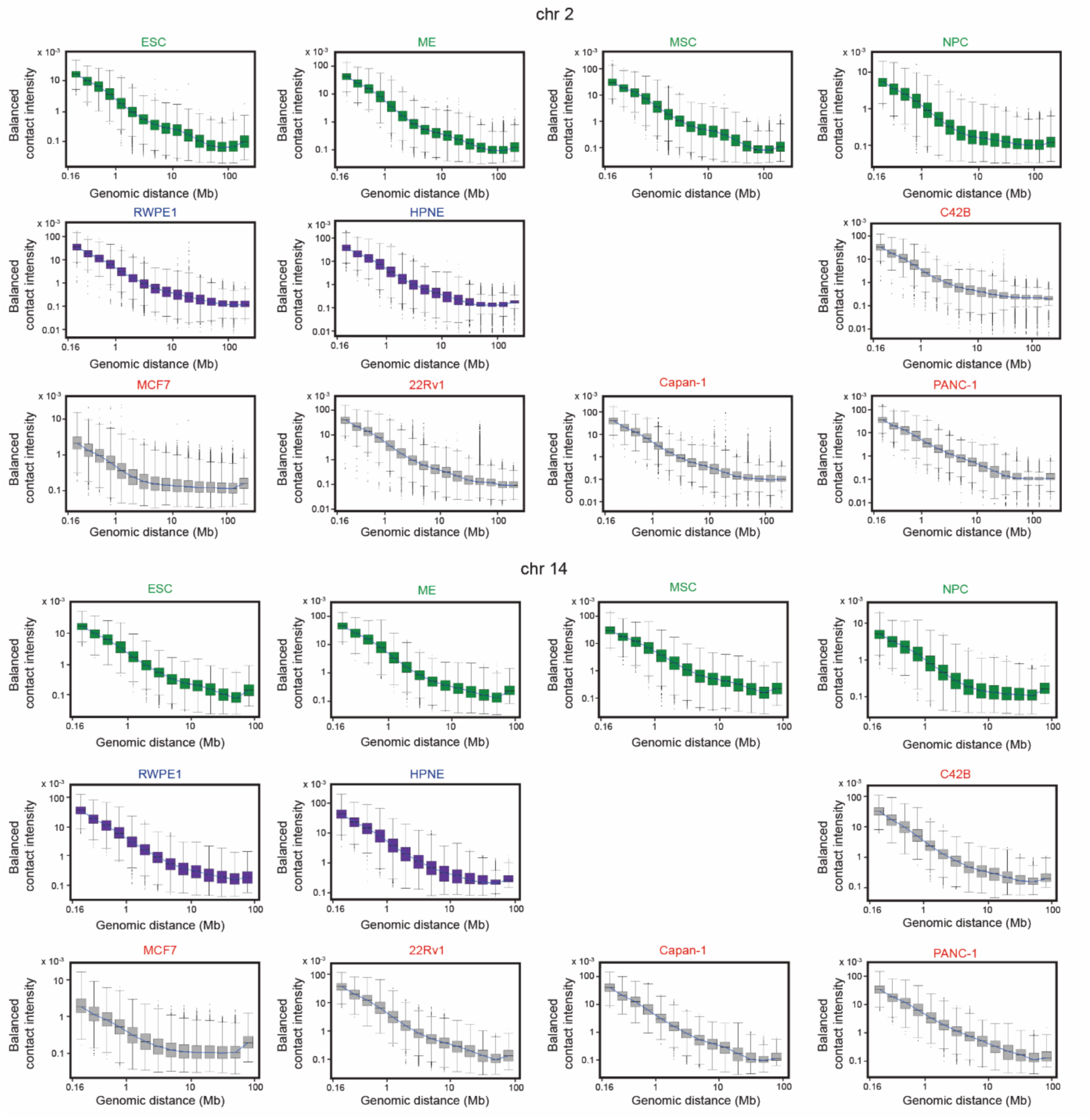
Prevalence of abnormally high contact frequencies in additional Hi-C datasets. Box plots of 11 additional Hi-C datasets show the widespread presence of genomic locus pairs separated by > 50 Mb in chr2 and chr14, exhibiting abnormally high contact frequencies.

**Supplementary Fig. S3.**
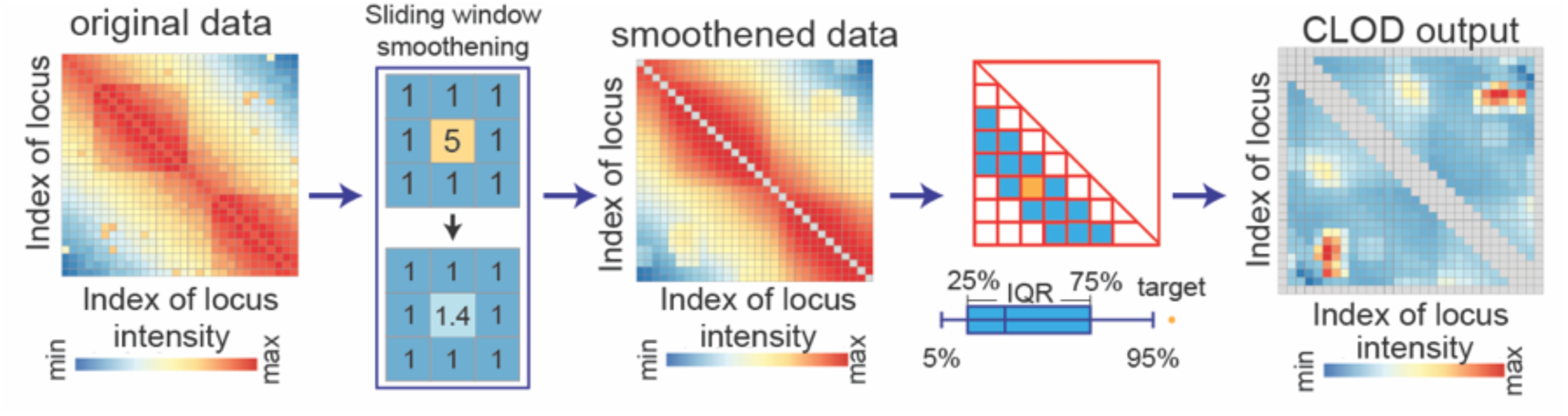
Chromosome Long-range colocalization identifier through Outlier Detection (CLOD) pipeline.

**Supplementary Fig. S4.**
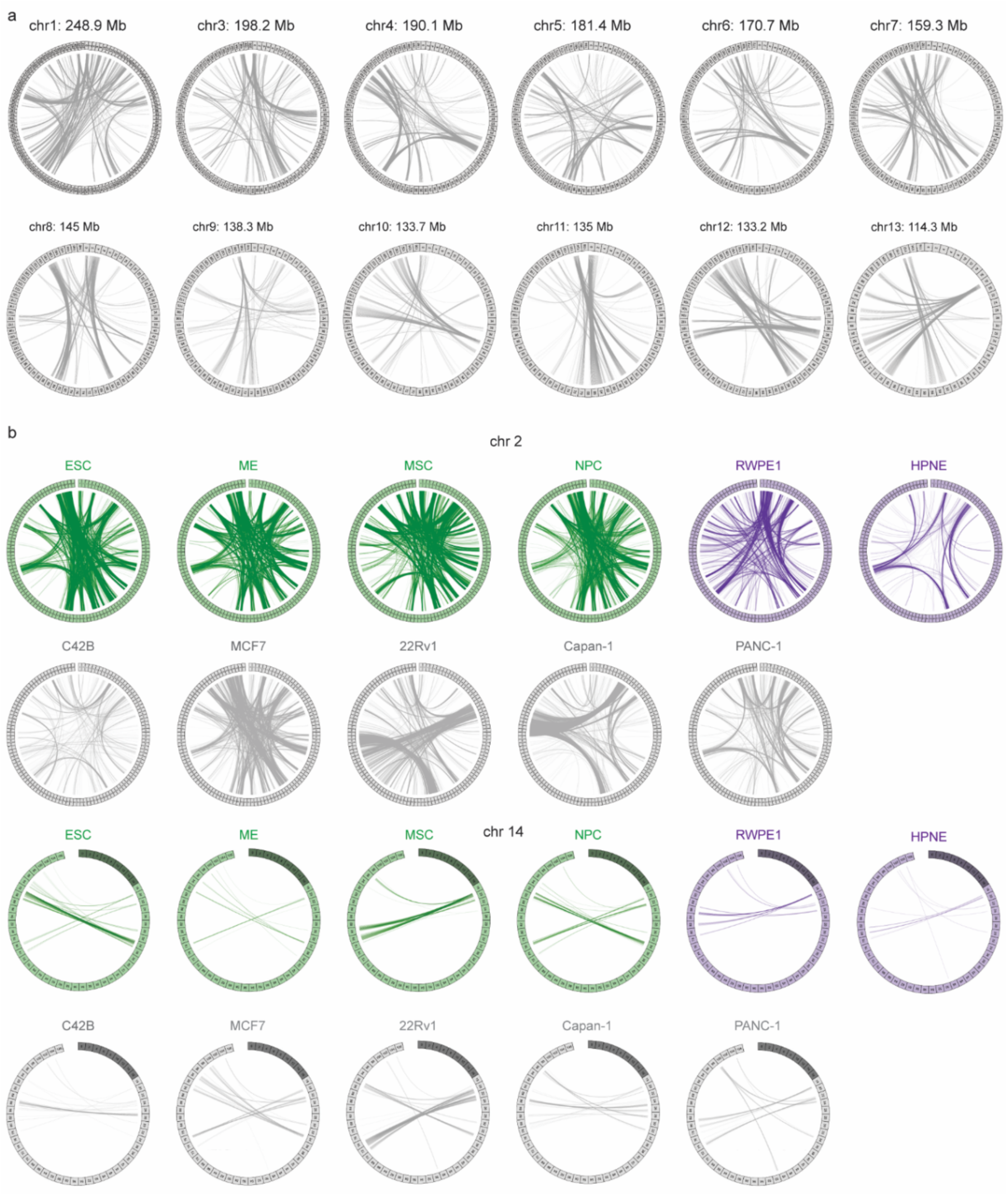
Chromosome linkage maps of CLOD-identified long-range colocalization loci. (**a**) Chromosome linkage maps depict CLOD-identified genomic locus pairs (>50 Mb apart) on chromosomes > 100 Mb in IMR90. (**b**) Chromosome linkage maps depict CLOD-identified genomic locus pairs (>50 Mb apart) in chr2 and chr14 that exhibit abnormally high contact frequencies across 11 additional cell lines.

**Supplementary Fig. S5.**
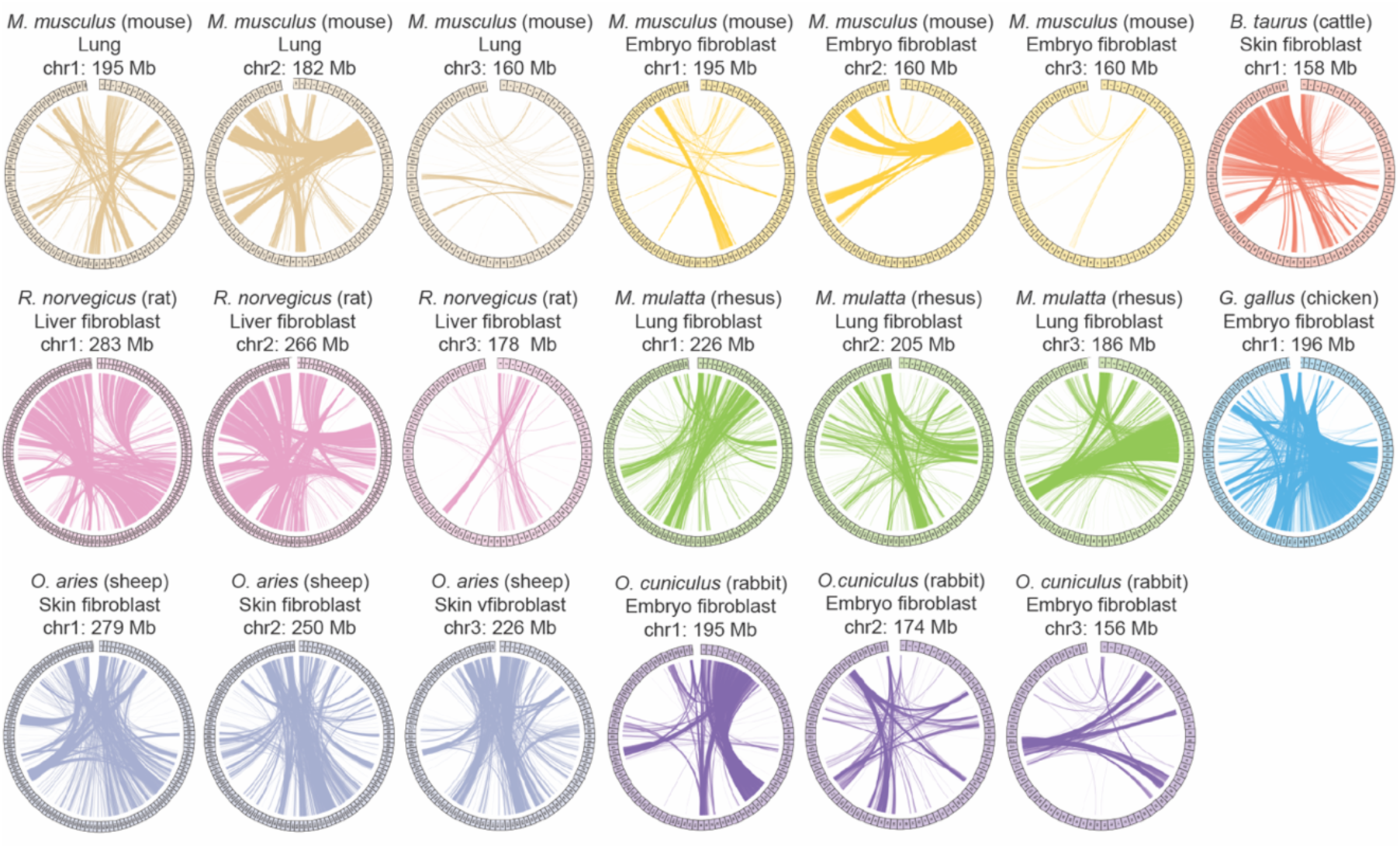
Chromosome linkage maps of CLOD-identified long-range colocalization loci from other species. Chromosome linkage maps depict CLOD-identified genomic locus pairs (>50 Mb apart) in six mammalian species and chicken.

**Supplementary Fig. S6.**
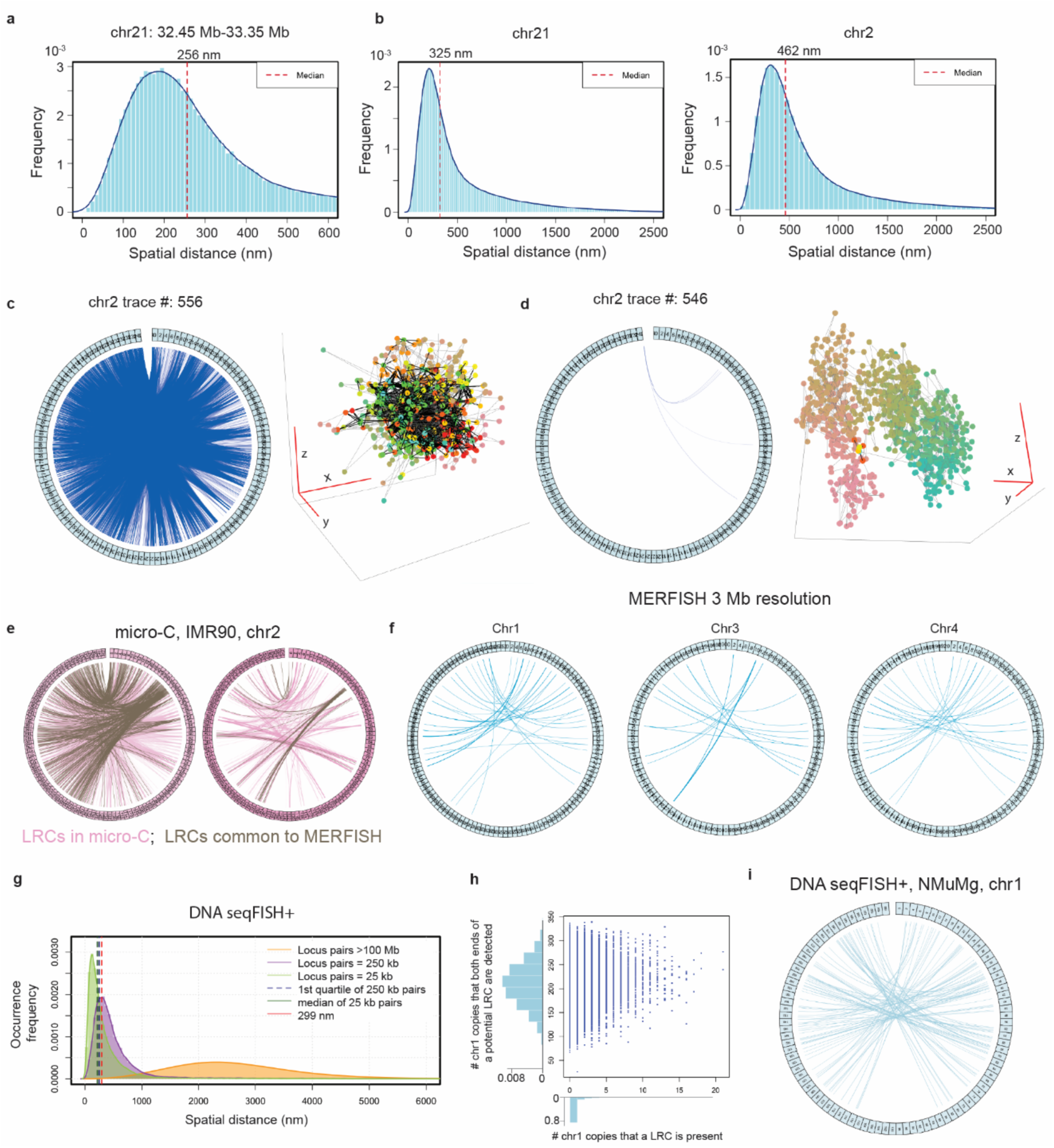
Statistical analysis and representative examples of individual chr2 traces from MERFISH chromatin tracing data. (**a**) Histogram and density curve illustrating the 3D distance distribution of adjacent labeled loci in chr21:32.45-33.35 Mb with the red dashed line indicating the median distance. (**b**) Histogram and density curves showing 3D distance distributions of adjacent labeled loci in chr21 (left) and chr2 (right), with red dashed lines representing the median distance between adjacent bins for each chromatin. (**c-d**) Linkage maps of LRCs and 3D structures of representative chr2 traces, where black lines denote LRCs and gray lines indicate neighbor links. (**e**) Linkage maps showing the long-range colocalization that identified with loose standard (left) and strict standard (right). Pink links are those identified from micro-C only, and the grey links are those also identified from MERFISH. (**f**) Linkage maps depicting sLRCs identified from other chromosomes in human cells from MERFISH data with 3 Mb resolution. (**g**) Histogram showing the distributions of spatial distances from pairs of loci with 25 kb (green), 250 kb (purple), or 100 Mb (orange) genome distance. (**h**) The scatter plot and histogram showing the number of chromatin that a certain pair of loci has been detected (y-axis) and the number of chromatin traces with the corresponding pair of loci colocalized (x-axis). (**i**) The linkage map showing the sLRCs in NMuMg chr1 from DNA seqFISH+.

**Supplementary Fig. S7.**
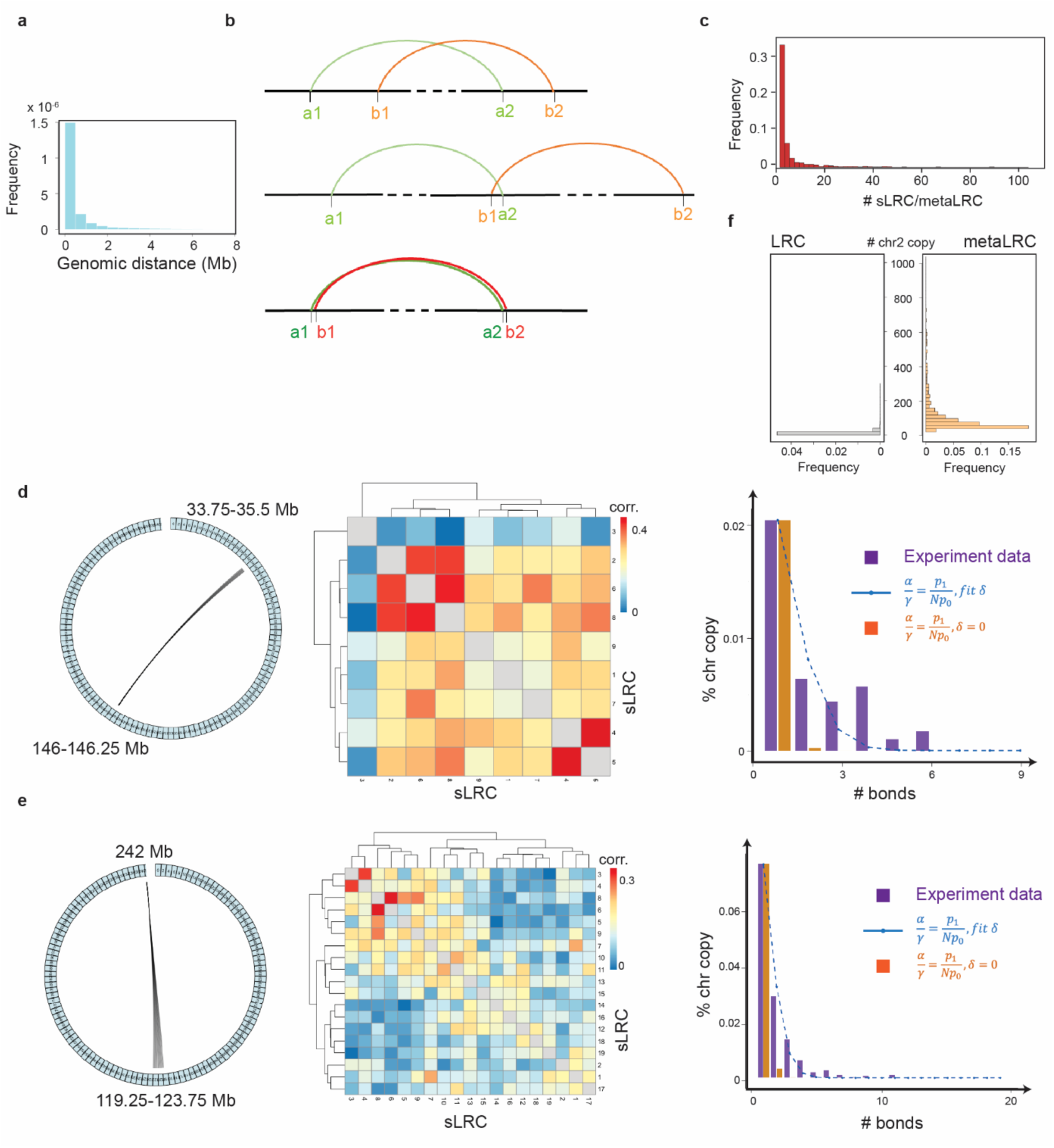
Additional results for multivalence binding model analyses. (**a**) Histogram depicting the distribution of genomic distances between two adjacent loci involved in sLRC. (**b**) Sketch plots illustrating three typical types of genomic relationships between sLRC a and sLRC b. (**c**) Density distribution of the number of sLRC pairs per metaLRC. (**d-e**) Linkage maps (left) of two representative metaLRCs, their heatmaps of correlation between sLRCs (middle), and probability comparisons (right) of observing various numbers of bound sLRCs within the metaLRC. The comparison includes experimentally observed frequencies (purple bars), predicted independent sLRC binding (orange bar), and the multivalent binding model (blue dashed line). *p_0_* is not shown. (**f**) Occurrence frequency of individual sLRCs (in gray) and metaLRCs (in orange) among the 2,991 traces of chr2.

**Supplementary Fig. S8.**
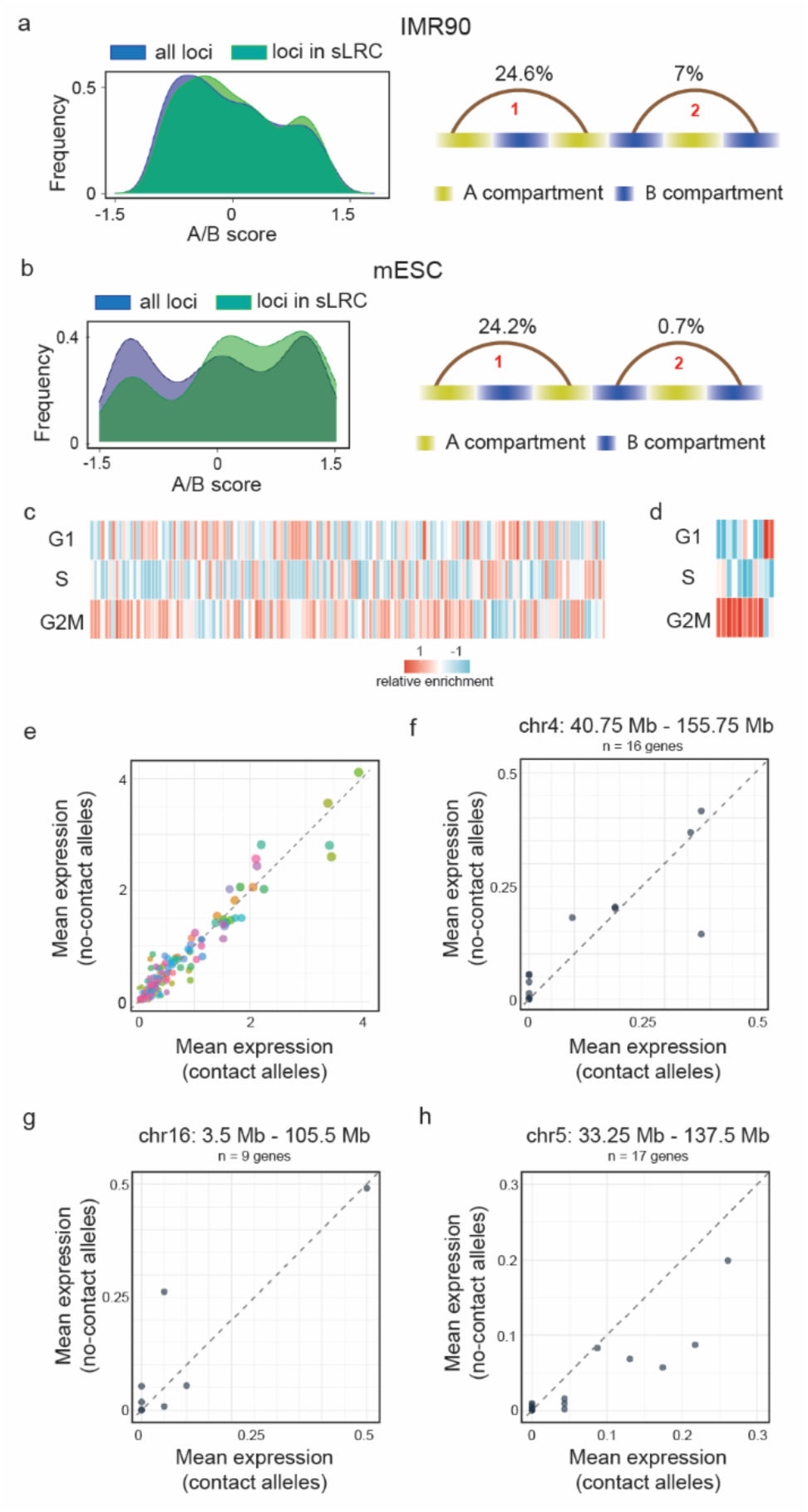
Association of long-range locus pairs connecting with cell cycle and gene activity. (**a-b**) Left: The distribution of A/B scores across all loci and only those associated with LRCs in IMR90 (a) and mESC (b), respectively. Right: percentage of LRCs with both ends in the same A/B compartment. (**c**) Cell cycle phase distribution of all LRCs. Each bar represents a locus pair, and the color intensity indicates the proportion of contact-forming alleles in each phase. (**d**) Pairs of loci that were significantly enriched in specific cell cycle phases. The bar and color are the same as panel c. (**e**) Comparison of mean gene expression at the anchors of all tested locus pairs between contact and no-contact alleles. Every point represents a gene, and genes located near the ends of the same locus pair share the same color. More details are present in the main text and Methods. (**f-h**) Mean expression of genes at the 40.75 Mb and 155.75 Mb loci on chr 4 (**f**), at the 3.5 Mb and 105.5 Mb on chr16 (**g**), and at the 33.25 Mb and 137.5 Mb on chr5 (**h**), respectively, comparing contact vs. no-contact alleles. Every point represents a gene.

**Supplementary Fig. S9.**
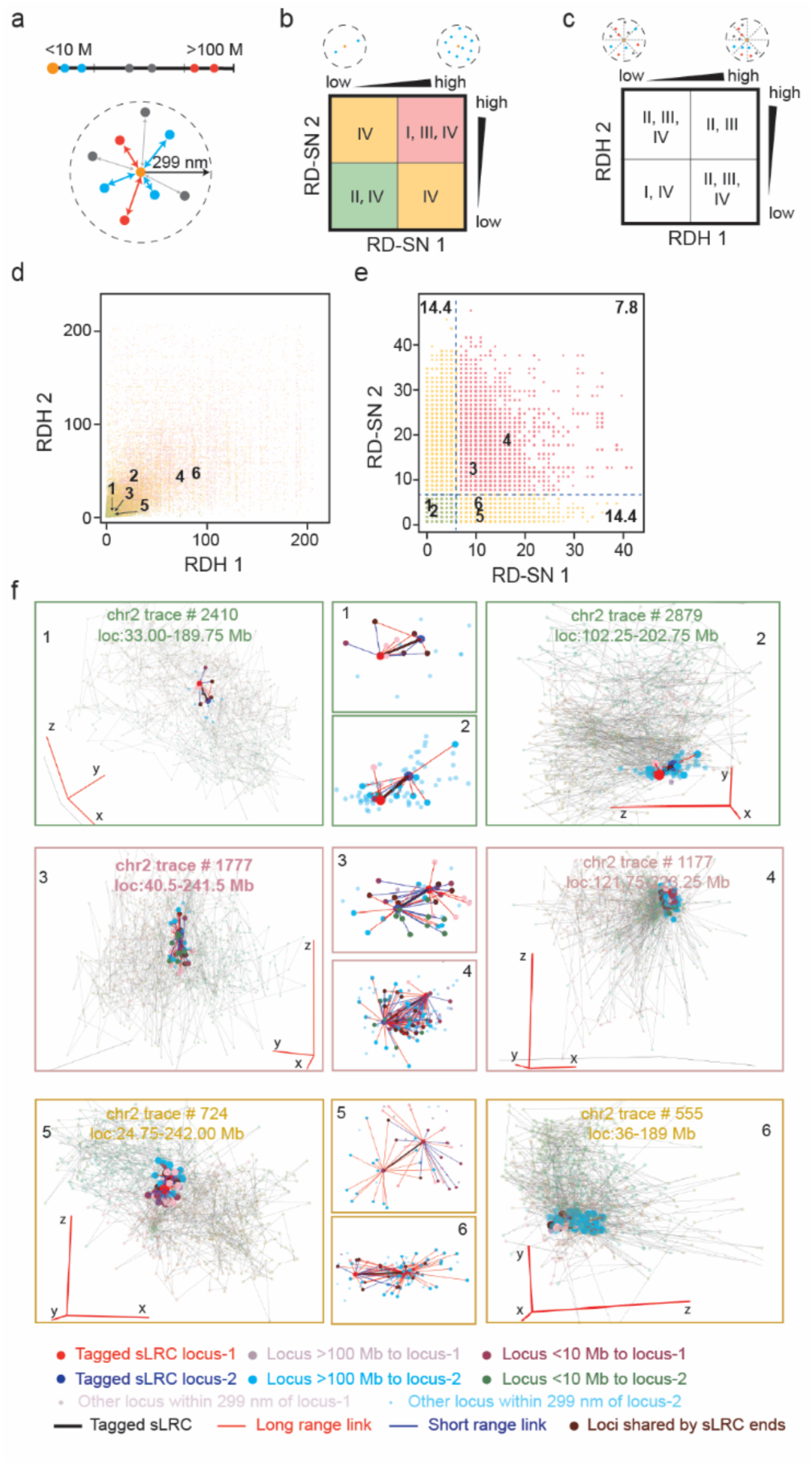
Possible mechanisms of sLRC formation and functional roles on chromosome folding. (**a**) Definition of the region around each locus from MERFISH data. (**b**) The diagram depicts the RD-SN defined in this paper. (**c**) The diagram depicts the RDH defined in this paper. (**d, e**) Scatter plots of RD-SN and RDH around both ends of individual sLRCs in each chr2 trace, respectively. Each dot represents one sLRC in a single chr2 trace. (**f**) 3D structures of six representative chr2 traces highlighted in panel d&e.

